# Copper Driven Mutualism of *Candida albicans* and *Staphylococcus aureus* Interkingdom Biofilms

**DOI:** 10.1101/2025.09.02.673504

**Authors:** Iana Kalinina, Roberto Vazquez-Muñoz, Orlando Ross, Philip A. Lewis, Kate Heesom, Philip Mitchelmore, Christian Hacker, Seána Duggan

## Abstract

Although fungi and bacteria commonly coexist within polymicrobial communities, the molecular mechanisms underlying their interactions are still not well understood. Here, we show that the fungus *Candida albicans* forms biofilms with the bacterium *Staphylococcus aureus* along a nutritional axis of mutualism and propose that “a copper economy” shapes fungal-bacterial biofilm interactions.

Using in vitro biofilms formed on plastic, we found that dual species biofilms are consistently larger than single-species counterparts, indicating a cooperative interaction. Dual species proteomic analysis revealed non-reciprocal copper handling: *C. albicans* increased copper uptake via transporter Ctr1, while *S. aureus* enhanced copper export via regulator CsoR and export chaperone CopZ. Dual species biofilms exhibited specific sensitivity to both copper depletion and supplementation, with corresponding reductions in biomass. We identified fungal copper import as the crucial element in mutualistic interactions between *C. albicans* and staphylococcal species. Moreover, fungal hyphae served as a critical scaffold for biofilm architecture, a role that was compromised under copper-replete conditions. Notably, copper nanoparticles disrupted these dual species biofilms, highlighting a potential therapeutic avenue. Furthermore, we extend the role of *C. albicans* copper import to mutualistic interactions with additional bacterial species. Our findings establish copper as a central mediator of *C. albicans* and *S. aureus* cooperation and suggest that a "copper economy" underpins mutualistic interactions in biofilms.

## Introduction

Fungi and bacteria co-inhabit diverse niches including soil, food and the human host, where they can co-exist as members of a healthy microbiome or co-infect to cause disease. *Candida albicans* is the most commonly isolated fungus from mixed fungal–bacterial infections (1), while *Staphylococcus aureus* is among the most frequently isolated bacteria from such infections (2). Individually, *C. albicans* and *S. aureus* are amongst the leading causes of human infection ranging from benign to life-threatening, world-wide (3,4). *C. albicans* and *S. aureus* occupy several of the same anatomical niches where they interact during both health and disease (5,6). The importance of this fungal-bacterial pairing is evident early in life, where competitive interactions lead to *C. albicans* inhibiting *S. aureus*, thereby shaping the infant gut microbiome (7). Later, there is the sustained simultaneous presence of both organisms in various microbiomes (8)(9) and finally, during disease both have been co-isolated from wound, oral, and systemic infections (10,11,12,13,14,15). Importantly, co-culture of both organisms from the blood stream is associated with a 50% mortality rate (16).

The interaction of *C. albicans* and *S. aureus* during in vitro and murine in vivo growth is described in the literature; as a bacterium, *S. aureus* sheds peptidoglycan fragments from its cell wall which triggers bacterial biofilm formation (17) and stimulates *C. albicans* to produce and elongate hyphae (18). *S. aureus* cells preferentially adhere to *C. albicans* hyphae via specific binding to the fungal adhesin molecules Als1/3 (19) and also non-specific binding (20,21). This physical interaction results in increased toxin production – particularly α-toxin – by *S. aureus,* via upregulation of the accessory genome regulator (Agr) (22). Furthermore, fungal ribose utilisation creates a low-ribose environment ensuring expression of *S. aureus* PurR and Agr which are required for purine biosynthesis and staphylococcal virulence in the context of intraabdominal infection (23).

Biofilms are aggregates of microbes which produce a polysaccharide matrix (24) and *C. albicans* and *S. aureus* cells readily interact to form biofilm, which is problematic as this mode of growth leads to enhanced antimicrobial resistance, frustrated clearance by cellular immunity and complicated treatment regimens (25,26). In dual species biofilms, *S. aureus* gains enhanced protection against vancomycin through *C. albicans* extracellular matrix (ECM) polysaccharides, farnesol production and biofilm regulator Bcr1 expression, and further benefits from facilitated translocation due to epithelial integrity disruption caused by *C. albicans* hyphae. (25,27,28). In turn, *C. albicans* also benefits from enhanced azole tolerance (11,29). Together, these interactions not only enhance the virulence and drug resistance of both organisms, but also pose a significant challenge for clinical management, underscoring the need to better understand and target the mechanisms underlying *C. albicans* and *S. aureus* dual species biofilm formation.

Mounting evidence supports that co-infections are common and result in worse disease outcomes, however there is a critical knowledge gap on the microbiological mechanistic basis of these infections (30). Improved understanding of fungal–bacterial interactions could reveal novel insights that enhance our ability to predict the outcomes of these interactions and inform strategies to control interkingdom interactions within complex microbial communities such as co-infections. Therefore, focusing on *C. albicans* and *S. aureus*, this study aimed to investigate the basis of mutualistic interaction in a mixed biofilm relevant to human disease.

## Results

### Characterisation of Mutualistic Biofilms

To develop a model of *C. albicans* and *S. aureus* biofilm, dual species biofilms were formed on plastic 96 well plates in RPMI-1640 media supplemented with 5% human serum and incubated at 37°C with 5% CO_2_. This condition, while firmly *in vitro*, included the requirement that microbial cells must adhere to plastic (as might happen on a catheter, voice prosthesis etc.) in the presence of RPMI-1640 and human serum, a mammalian cell culture medium loosely mimicking conditions in a human host. Equal fungal and bacterial cell numbers were allowed to adhere for 4 hours, and then non-adherent cells were removed by gentle washing. Media was replenished and biofilms then matured up to three days at 37°C and 5% CO_2_ (Fig 1a). Biofilm metabolic activity was measured via XTT assay (Figure 1b), and total biomass was determined via crystal violet assay (Figure 1c). Both approaches demonstrated biofilms increased biomass and metabolic activity over time, up to 72 hours. Fungal single species biofilms had greater biomass than bacterial single species biofilms across the time course, and dual species biofilms had consistently greater biomass than bacterial biofilms throughout the experiment. At 48 hours, the dual species biofilms had significantly greater biomass than either of its single species counterparts (*C. albicans* versus dual species *P value* =0.007 and *S. aureus* versus dual species *P value* =0.0001). We proceeded to investigate the dual species biofilms at this time point as it consistently provided evidence of a mutualistic interaction. We defined mutualistic as a readout in the dual species condition greater than the expected readout for the single species condition. Next, we visualised the biofilms using scanning electron microscopy (SEM) and observed distinct features in each biofilm. *C. albicans* biofilms consisted of both yeast and hyphal structures and some matrix production was observed (Fig 1d). *S. aureus* biofilms appeared as a film of uniform spherical cells and minor matrix production was observed (Fig 1e), likely due to the inhibitory effect of human serum on *S. aureus* biofilm formation and matrix production in vitro (31,20). Dual species biofilms showed physical interaction of *C. albicans* and *S. aureus*, with an enrichment of bacterial cells at hyphal structures (Fig1f), in line with previous reports (19,25, 22).

**Figure 1.**
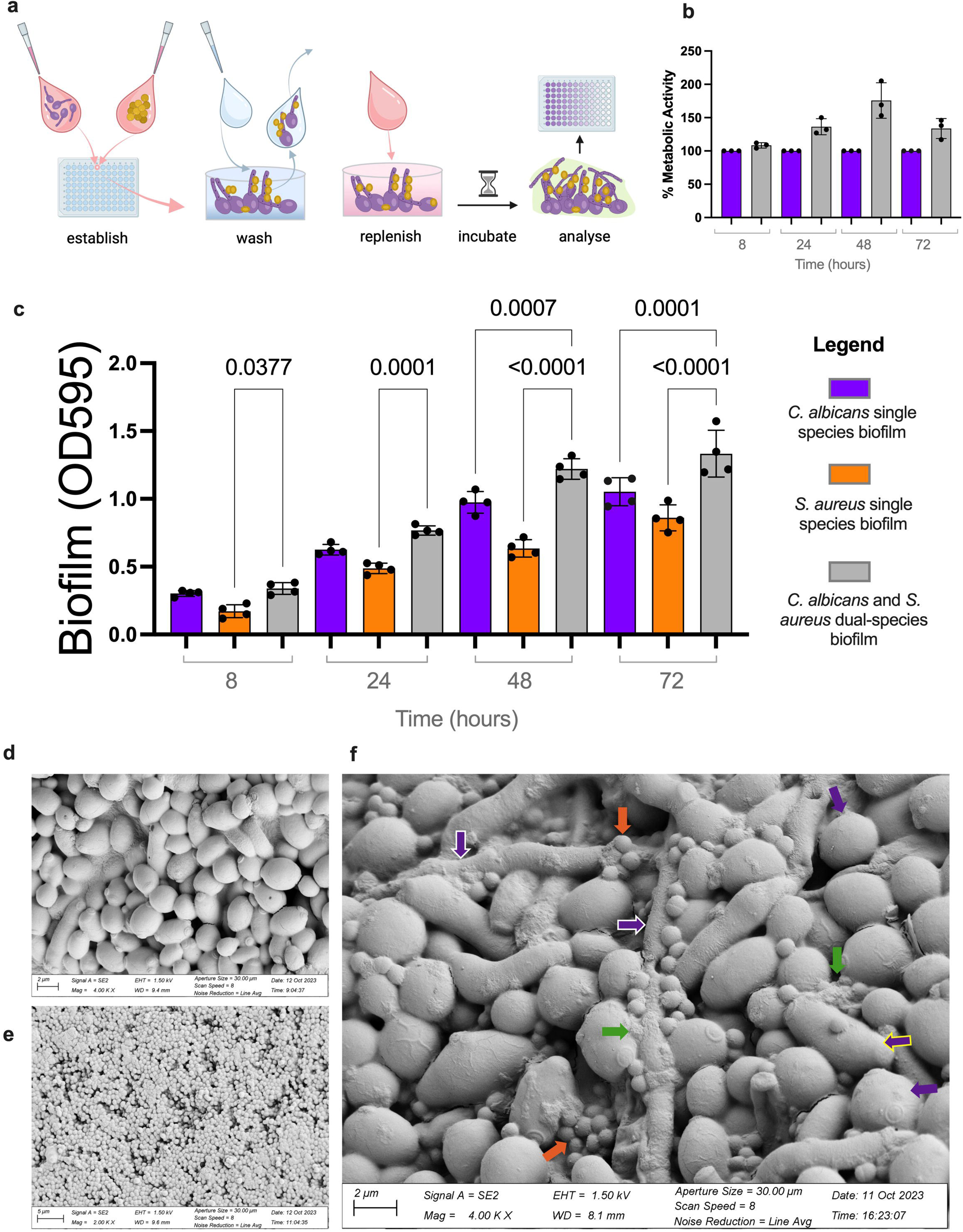
Characterisation of Mutualistic *C. albicans* and *S. aureus* Biofilms. **a** Dual species biofilms of *C. albicans* and *S. aureus* were established in RPMI containing 5% human serum. Non-adherent cells were washed away, and biofilms were cultured to maturity before further analysis. **b** Metabolic activity of total biofilm was determined at 8, 24, 48 and 72 hours after washing via XTT assay. Dual species biofilm metabolic activity (shown in grey) was normalised to single species fungal metabolic activity. Data points represent individual biological replicates. Data shown are mean and standard error of the mean (SEM). **c** Total biomass of single and dual species biofilms was determined via CV assay at 4, 8, 48 and 72 hours after washing. Data are shown as absolute absorbance values determined at optical density 595nm where each data point represents a biological replicate. Error bars represent standard deviation. Statistical significance was determined using a One-Way Anova with multiple comparisons and Sidak’s correction and P values are shown as numerical values. **d** Representative scanning Electron Microscopy (SEM) of 48 hour *C. albicans* biofilm **e** Representative SEM of 48 hour *S. aureus* biofilm **f** Representative SEM of 48 hour dual species biofilm of *C. albicans* and *S. aureus* where fungal hyphae are indicated by purple arrows with white border, yeast cells indicated by purple arrows, elongated yeast cells indicated by purple arrows with yellow border, bacterial cells indicated by orange arrows and biofilm matrix indicated by green arrow.

### Dual Species Biofilm Proteomics

Investigating the basis of mutualistic pathogen interactions will be essential for understanding the mechanisms underpinning microbial cooperation and community stability both which contribute to outcomes in health and disease. Therefore, to identify the processes contributing to *C. albicans* and *S. aureus* mutualistic interaction during biofilm growth, we chose to investigate biofilms at 48 hours, which were significantly larger and more metabolically active than single species controls. This was mirrored in biofilm protein content, where dual species biofilms contained more protein at 48 hours (*C. albicans* versus dual species *P value* =0.104 and *S. aureus* versus dual-species *P* value =0.0013) (Fig 2a). Total protein was isolated from the single and dual species biofilms and an equal volume (100 μL) from each condition was subject to proteomics via Tandem Mass Tagging-Mass Spectroscopy (TMT-MS). Our approach compared the proteome of single species biofilms to the respective species proteome in the dual species biofilm (Fig 2b). Principal component analysis (PCA) (Fig 2c) revealed that biofilm proteomes cluster according to species, with dual species biofilm clustering together. Biofilms are complex structures of viable and possibly non-viable cells, as well as matrix and cell debris, existing across gradients of nutrients, oxygen and density. In addition to this complexity, it appears each biofilm develops a distinct proteome. The individual biofilm variance hampers P values from being as low would be expected in single state, or planktonic cultures. Nevertheless, our approach detected over 4000 proteins. A two tailed, equal variance *t*-test with a *P* value of <0.05 and a fold change of >2 cut-off resulted in a list of 329 *C. albicans* proteins and 277 *S. aureus* proteins. Proteins meeting these criteria were subject to functional classification according to protein class by the PANTHER database (32) (Fig 2d and Fig 2f). Despite distinct *C. albicans* and *S. aureus* classification patterns generally, metabolic interconversion enzyme was the highest represented class of both proteomes, followed by translational proteins. Volcano plots for *C. albicans* proteins show the most highly expressed and significant proteins in purple (Fig 2e) while significant proteins for *S. aureus* are show in orange (Fig 2g).

**Figure 2.**
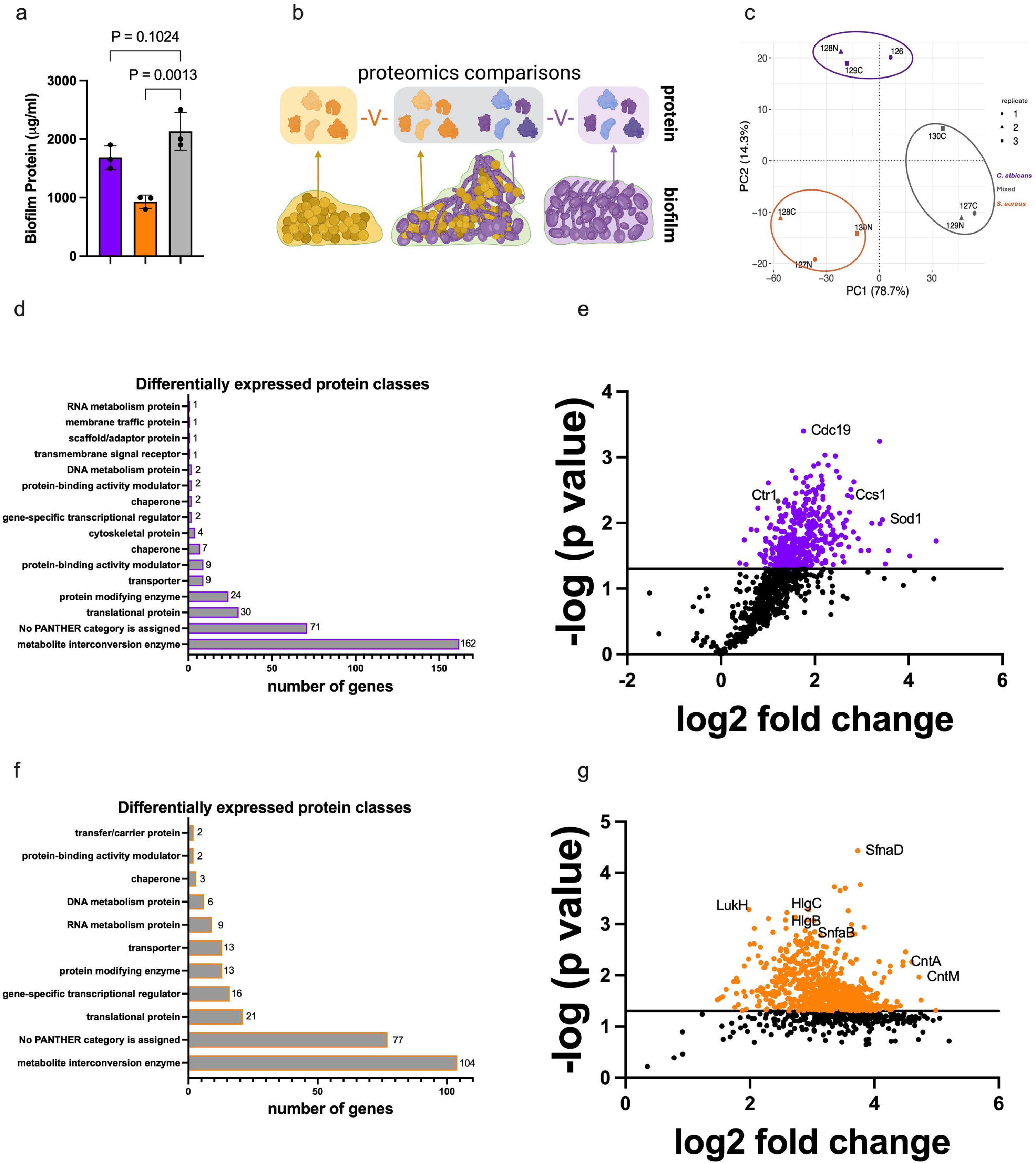
Comparative Proteomics of *C. albicans* and *S. aureus* in Mixed Biofilms. **a** Total protein was extracted from single and dual species biofilms and quantified via BCA assay. *C. albicans* biofilm protein is shown in purple; *S. aureus* biofilm protein is shown in orange and dual species biofilm protein is shown in grey. **b** A schematic demonstrating the proteomic analysis approach whereby *S. aureus* protein (shown in orange) from single species biofilms was compared to *S. aureus* protein from dual species biofilms, and *C. albicans* protein (shown in purple) from single species biofilms was compared to *C. albicans* protein from dual species biofilms. **c** Principal component analysis of all biofilm proteomes captured shows *C. albicans* single species biofilms in purple, *S. aureus* single species biofilms in Orange and dual species biofilms in grey. The data point shapes correspond to the replicate within the experiment n=3. **d** Quantification of the number of *C. albicans* proteins regulated in PANTHER protein classes. **e** Proteome comparison of *C. albicans* single species biofilm and *C. albicans* within a dual species biofilm with *S. aureus*. Only *C. albicans* proteins are shown. Proteins with absolute fold changes of >2 and *P* value <0.05 are considered to be differentially expressed and included in further analysis. Proteins were assigned identifiers using annotations from UNIPROT. **f** Quantification of the number of *S. aureus* proteins regulated in PANTHER protein classes. **g** Proteome comparison of *S. aureus* single species biofilm and *S. aureus* within a dual species biofilm with *C. albicans*. Only *S. aureus* proteins are shown. Proteins with absolute fold changes of >2 and *P* value <0.05 are considered to be differentially expressed and included in further analysis. Proteins were assigned identifiers using annotations from UNIPROT.

### *C. albicans* proteomic response to mixed biofilm with *S. aureus*

Consistent with greater biomass and protein content observed at 48 hour biofilms in Fig 1c and Fig 1d, there is also increased expression of protein synthesis and translation proteins such as Tif5, Tif6, Anb1 and Efb1; Ribosomal subunits Rpps27, Rpp2B, Rps27A, Rps21B, Rps28B, Rps3, Rpp2A, Rps0, Rps10; Chaperones Ccs1, ChzI, Asf1; And heat shock proteins such as Hsp21, Hsp104, Ssb1, as well putative proteins involved in these pathways A0A1D8PHU2 and A0A1D8PFG6. Proteins involved in interaction with the human host and virulence are also up-regulated including Rhd3 (FC 7.07; *P* value 0.013), Sap9 (FC 2.8; *P* value 0.043) and superoxide dismutases (SODs - presented later). As expected, metabolism is impacted by mixed biofilm growth with increases in amino acid synthesis (Ser33), carbohydrate metabolism (Gph1, Eno1, Tdh3, Pgk1, Gor1) and lactate metabolism (Glo2, Aip2). The most differentially regulated *C. albicans* protein in response to *S. aureus* mixed biofilm growth is superoxide dismutase Sod1 (FC 10.85; *P* value 0.0089), an antioxidant which catalyses the dismutation of superoxide radical anions to oxygen and hydrogen peroxide (33). Growth with *S. aureus* in a dual species biofilm induces further oxidative stress response proteins such as the superoxide dismutase 1 copper chaperone Ccs1 (FC 6.49; *P value* 0.0038), thioredoxin peroxidase Prx1 (FC 3.68; *P* value 0.012) and superoxide dismutase Sod2 (FC 3.25; *P* value 0.028). This profile demonstrates that dual species biofilm growth with *S. aureus* is an oxidative stress condition for *C. albicans* and is in agreement with the data from Peters and colleagues who speculated that the presence of *S. aureus* induced a stress response by *C. albicans* in dual species biofilm (34).

### *S. aureus* adaptation to *C. albicans* mixed biofilm

Mirroring its fungal partner and consistent with increased protein content and larger biomass, *S. aureus* increases expression of the chaperone activity with GroES (FC 6.61 *P* value 0.02) and GroEL (FC 17.2; *P* value 0.061). The GroES/GroEL system assembles into a nano-cage that encapsulates non-native substrate proteins, creating a specialized physical environment that facilitates and accelerates their proper folding. Additionally, Clp proteins ClpL (FC 12.99; *P* value 0.068), ClpB (FC 5.3; *P* value 0.04), ClpC (FC 7.3; *P* value 0.022) and ClpP (FC 15.9; *P* value 0.05) are increased providing further proteolytic and chaperone machinery for bacterial protein synthesis. Expression of Clp family proteins is not surprising as these are required for stress tolerance and biofilm formation (35) – both of which our fungal-bacterial model provide. Interestingly, *S. aureus* increases expression of repressor of toxins Rot (FC 19.07; *P* value 0.006) and SarS (FC 8.11; *P* value 0.006), both of which are repressors of the accessary genome regulator (Agr) responsible for quorum sensing, Agr regulated toxicity and general virulence in *S. aureus*, thereby indicating a suppression of classical virulence traits while in biofilm growth with *C. albicans*. However, we did detect upregulation of the core-genome encoded leukocidin Gamma-Hemolysin CB HlgC (FC 6.03; *P* value 0.0005) and HlgB (FC 5.9; *P* value 0.0008). Given the phenol soluble modulin toxins harbour anti-*Candida* activity (36), it’s tempting to speculate that bacterial toxin expression could be orchestrated to manage the interaction with *C. albicans* within the biofilm.

Increased Sortase A SrtA expression (FC 7.6; *P value* 0.036) permits surface protein anchorage on the cell wall, such as virulence factors (37). *S. aureus* biofilm matrix is largely composed of DNA, which is released via the hydrolytic activity of autolysin Atl (FC 3.7; *P* value 0.02). Building on this effort to construct a matrix, Extracellular matrix-binding protein Ebh is also up-regulated (FC 10.38; *P* value 0.08). In mixed biofilms with *C. albicans*, *S. aureus* benefits from enhanced antimicrobial tolerance (25). In our data, we detect increases not only in matrix production but also response to drugs via up-regulation of penicillin binding protein 1 Pbp1 (FC 5.75; *P* value 0.004), Ppb2 (FC 7.4; *P* value 0.04), Pbp4 (FC 7.8; *P* value 0.03), Pbp3 (FC 4; *P* value 0.002) and VraR (FC 7.9; *P* value 0.008). Further mirroring the proteomic response to mixed biofilm growth of *C. albicans*, *S. aureus* also strongly regulated a protein putatively involved with detoxification, the DoxX protein Q2G2G1 (FC 17; *P* value 0.06) as well as thioredoxin TrxA (FC 7.3; *P* value 0.013), and putative thioredoxins Q2FWX1(FC 6.8; *P* value 0.01), Q2G000 (FC 7.8; *P* value 0.016) and Q2FWB6 (FC 11.9; *P* value 0.035). The *S. aureus* mixed biofilm proteome is characterised by a concerted shift in purine biosynthesis (PurQ, PurN, PurK, PurF, PurD and DeoD are all significantly upregulated, with many more proteins in the pathway expressed, but below our *P value* cut-off). Purines are essential aromatic macromolecules, with their importance underscored by the capacity of microbes to produce them *de novo*. Purine synthesis and regulation via PurR are required for *S. aureus* virulence and nutrient acquisition in the host (38) and also during co-infection with *C. albicans* (23). Our data broaden their importance to interaction with *C. albicans* in biofilms. Lastly, we observed a broad effort to acquire metals in *S. aureus*; siderophore activity is increased by SfnaD (FC 13.3; *P* value 0.0004) and SfnaB (FC 7.8; *P* value 0.001), while the broad spectrum metallophore staphylopine is increased via CntA (FC 22; *P* value 0.005) and CntM expression (FC 26.3; *P* value 0.01). This prompted a closer examination of the roles of metals.

### A role for copper in *C. albicans* and *S. aureus* mixed biofilms

We noticed that processes relating to copper transport were affected in both *C. albicans* and *S. aureus* in mixed biofilms. In dual species biofilms *C. albicans* produced more Copper transport protein 1 Ctr1, Copper binding protein Sco1, Copper requiring Superoxide dismutase 1 Sod1 and Sod1 copper chaperone Ccs1 (Table 1). Meanwhile, *S. aureus* increased expression of Copper chaperone CopZ and Copper sensing transcriptional repressor CsoR (Table 1). *C. albicans* Ctr1 is a transmembrane copper ion importer required for high affinity copper transport into the cell. It is expressed under copper-limiting conditions and is positively regulated by the transcription factor Mac1, which was previously shown to be differentially regulated in mixed biofilms with *S. aureus* (34). Ctr1 expression is induced during interaction with macrophages, formation of biofilm, and in response to alkaline pH via the Rim101 pathway. When disrupted Ctr1 mutants grow poorly on low copper solid medium and have altered morphology in response to copper deletion (39,40) (41)(42). *S. aureus* CopZ is a copper chaperone that binds and delivers copper ions to the exporter CopA, with its expression induced by elevated copper levels to help reduce intracellular copper concentration (43). These data indicate that within the dual species biofilm, *C. albicans* increases import of copper ions and we hypothesise this may function to support activity of Sod1. Meanwhile, *S. aureus* increases expression of CopZ which shuttles copper to CopA for export, and a repressor of copper machinery induced by high copper, CsoR. Together, this suggests *C. albicans* acquires copper while *S. aureus* exports it. It is tempting to speculate at this point that within the mixed biofilm the fungus and bacterium collaborate to maintain “copper stasis” which supports Sod1 expression which in turn is important for the fungal stress response during interaction with bacteria. As this activity is present during mutualistic biofilm growth, we hypothesised that this copper response was a trait important for mutualism.

**Table 1.** Copper Protein Expression by *C. albicans* and *S. aureus* in Dual Species Biofilms. Protein name, protein function, species, absolute fold change (FC) and *P value* are listed for proteins related to copper processed by both *C. albicans* and *S. aureus*. Name and function were determined by cross referencing accession identifier with data in UniProt. The FC was calculated for three biological replicates by averaging the raw protein abundance and dividing the dual species biofilm raw abundances by the single species biofilm raw abundances. *P* value was determined using a two-sided Student’s *t*-test of equal variance.

| Protein Name | Protein Function | Species | FC | P value |
| --- | --- | --- | --- | --- |
| Sod1 | Superoxide dismutase [Cu/Zn] | *C. albicans* | 10.85 | 0.0089 |
| Ccs1 | Superoxide dismutase 1 copper chaperone | *C. albicans* | 6.49 | 0.0038 |
| Ctr1 | Copper transport protein | *C. albicans* | 2.56 | 0.004 |
| Sco1 | Copper binding protein | *C. albicans* | 2 | 0.018 |
| CsoR | Copper sensing transcriptional repressor | *S. aureus* | 9.65 | 0.045 |
| CopZ | Copper chaperone | *S. aureus* | 4.68 | 0.007 |

### Copper is an axis of Mutualism for *C. albicans* and *S. aureus* Biofilms

To determine if alterations in environmental copper impact single or dual species biofilms, total biofilm biomass was measured by crystal violet assay. Biofilms attached in standard conditions and were replenished with either standard medium, medium containing copper sulphate (CuSO_4_) (Fig 3a) or medium containing bathocuproine sulphonate (BCS – a copper-specific chelating agent) (Fig3b). Interestingly, the inhibitory CuSO_4_ concentrations for single species biofilms was higher than for dual species biofilms. *S. aureus* biofilms were inhibited at 0.01 mM (*P* value = 0.0033 compared to 0 mM CuSO4), whereas *C. albicans* biofilms were inhibited at 5 mM (*P value* = 0.0029 compared to 0 mM CuSO_4_). Biomass of dual species biofilms is 1.5-fold greater than the respective single species biofilms at 0 mM CuSO_4_ and these biofilms are deemed mutualistic. Mutualistic biofilm formation was lost at 0.05 mM CuSO_4_ (*P* value = 0.0001 compared to 0 mM CuSO_4_). Therefore, at 0.05 mM CuSO_4_ single species biofilms were indistinguishable from control conditions, while dual species biofilms failed to reach the biomass observed in standard conditions. This indicated that mutualism did not occur at this concentration of copper. Similarly, when copper was chelated from the media at 1 µM BCS single species biofilms were unaffected, but dual species biofilms no longer reached their mutualistic biomass levels. Propensity to form a mutualistic biofilm was calculated in CuSO_4_ (Fig 3c) and BCS (Fig3d). To investigate if our observation was copper specific, or rather a general metal ion effect, magnesium and zinc were included in the culture medium (Fig 3e and Fig3f). Single and dual species biofilms were impacted similarly, i.e., there was no dual species-specific effect, in increasing magnesium and zinc concentrations up to 300 μM, which was the range within which we observed the effect of copper. To determine the effect of the environmental copper conditions pertinent to biofilm mutualism on planktonic cultures, we also performed growth curves in standard liquid media compared to copper replete and deplete media (Supplementary Figure 1). The copper replete (50 μM CuSO_4_) and deplete (1μM BCS) conditions at which mutualism was lost during biofilm growth, but at which single species biofilms were not affected impact the growth of planktonic cultures. *C. albicans* was slightly delayed reaching log phase when copper was chelated, while *S. aureus* failed to grow in 50 μM CuS04 in liquid. This indicates *C. albicans* and *S. aureus* harbour copper resilience in biofilm, which is not evident in the planktonic mode of growth at the concentrations tested. We therefore proceeded to explore the role of copper in mutualistic *C. albicans* and *S. aureus* biofilms.

**Figure 3.**
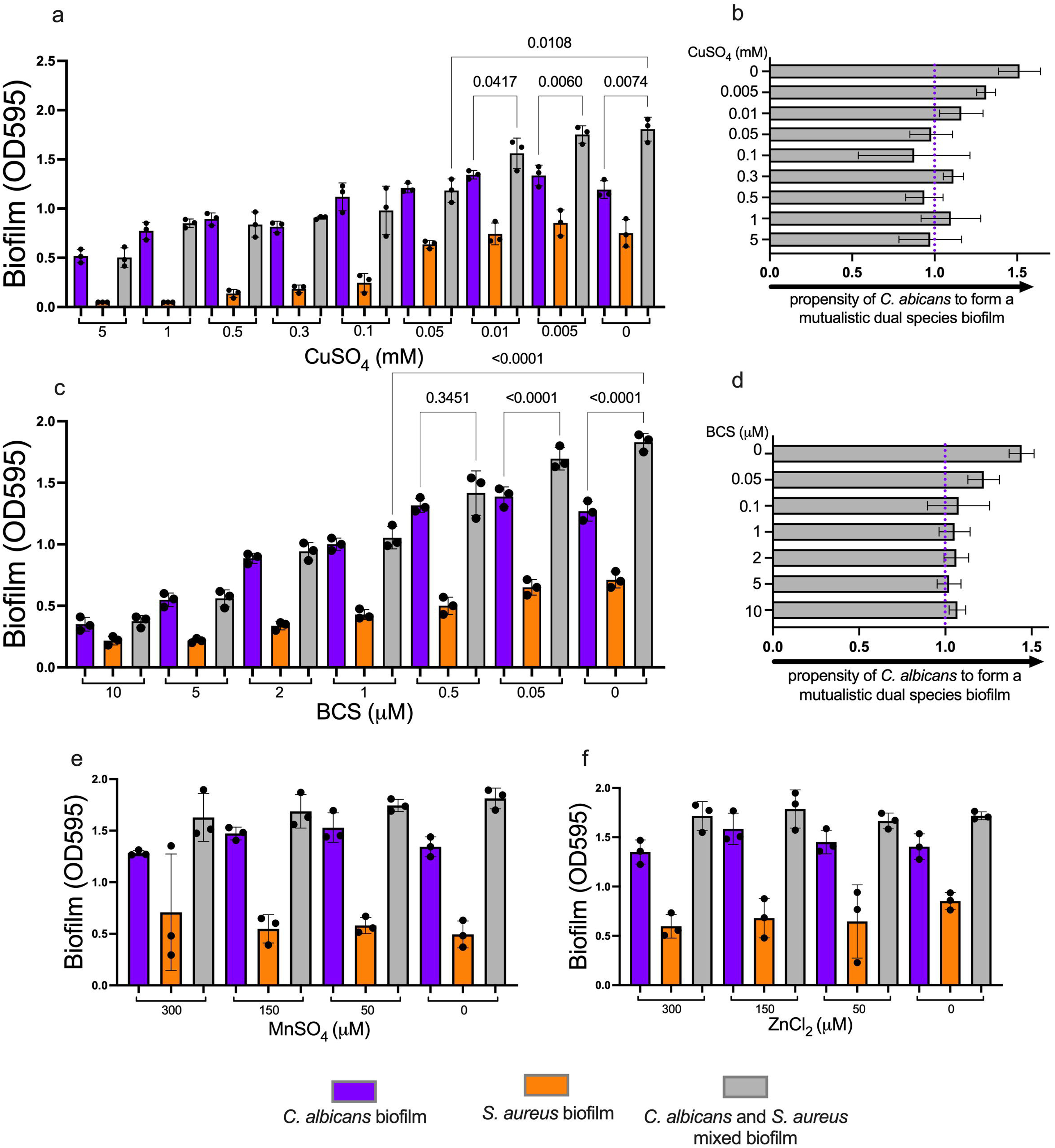
Copper is an axis of Mutualism for *C. albicans* and *S. aureus* Biofilms. **a** Total biomass of single and dual species biofilms was determined via CV assay after 48 hours of biofilm culture in various concentrations of CuSO_4_ranging from 0 – 5 mM. Data are shown as absolute absorbance values determined at optical density 595nm where each data point represents a biological replicate. Error bars represent standard deviation. Statistical significance was determined using a One-Way Anova with multiple comparisons and Sidak’s correction and P values are shown as numerical values. **b** The propensity of *C. albicans* to form a mutualistic dual species biofilm with *S. aureus* in various concentrations of CuSO_4_ was determined by comparing the total biomass of single species fungal biofilm to the biomass of dual species biofilms to generate a value representing the propensity to form mutualistic dual species biofilms with *S. aureus* (OD595 of dual species biofilm / OD595 single species biofilm = propensity to form a mutualistic biofilm) whereby a value of 1 (indicated by a dashed purple line) indicates a biofilm the same as the single species biofilm, a value less than 1 indicates an antagonistic biofilm and a value greater than 1 indicates a mutualistic biofilm. This calculation was performed for all concentrations of CuSO_4_ using the data from (**a)**. Data are represented as grey bars showing the mean and errors bars indicate standard deviation. **c** Total biomass of single and dual species biofilms was determined via CV assay after 48 hours of biofilm culture in various concentrations of the copper-specific chelator BCS ranging from 0 – 10 µM. **d** The propensity to form a dual species biofilm in copper deplete conditions is shown. Values calculated as in (**c)**. **e** Total biomass of single and dual species biofilms was determined via CV assay after 48 hours of biofilm culture in various concentrations of MnSO_4_ ranging from 0 – 300 µM. Neither single species nor dual species biofilm was affected at these concentrations. Data are shown as absolute absorbance values determined at optical density 595nm where each data point represents a biological replicate. Error bars represent standard deviation. **f** Total biomass of single and dual species biofilms was determined via CV assay after 48 hours of biofilm culture in various concentrations of ZnCl_2_ ranging from 0 – 300 µM. Neither single species nor dual species biofilm was affected at these concentrations. Data are shown as absolute absorbance values determined at optical density 595nm where each data point represents a biological replicate. Error bars represent standard deviation.

### Environmental Copper Effects Dual Species Biofilm Architecture

As we established that mutualistic biofilm biomass was influenced along an axis of copper, we wondered how the physical structure of the biofilm was impacted. To capture structural changes fungal-bacterial interaction within the biofilm, we cultured dual species biofilms on standard (RPMI), copper replete (50 μM CuSO_4_) or copper deplete (1 μM BCS) agar pads – a method to permit imaging via scanning electron microscopy (Fig 4a-c). Prior to imaging, we noticed the biofilms appeared to have distinct structures with the naked eye (Fig 4d). Control biofilms of *C. albicans* appeared to have protrusions around the ring of the colony and a ruffled centre, consistent with hyphal morphology, while *S. aureus* grew as a creamy-brown, smooth colony. Dual species biofilm grew larger than either single species and appeared to have a pigmented centre and a wrinkled topology (Fig 4d). Biofilms cultured in the copper replete condition were visibly different from the controls. *C. albicans* did not produce protrusions or ruffling, and the dual species biofilm was of uniform pigmentation (with the naked eye) and had a smooth topology (Fig 4d). Notably, *S. aureus* appeared more pigmented in the copper replete condition with a deeper brown colour enriched at the centre of the colony. The loss of fungal morphology was mirrored in our electron microscopy data. Biofilms grown on CuSO_4_ (Fig 4b) demonstrated a remarkable loss of hyphal structure compared to control (Fig 4a) or copper deplete conditions (Fig 4c). Copper deplete biofilms were indistinguishable from controls. The presence of yeast or hyphal *C. albicans* morphotypes profoundly effects architecture as the biofilm matures (44) and we hypothesise that this loss of hyphal structure underpins the loss of mutualism observed in copper replete conditions. To determine if the fungal-bacterial make-up of the biofilm was also affected by copper, we quantified *C. albicans* and *S. aureus* colony forming units (CFU) on selective media. In standard conditions, the proportion of fungi to bacteria determined by CFU is approx. 35% *C. albicans* and 65 % *S. aureus* (Fig 4e). However, when environmental copper is increased or decreased, these proportions shift, resulting in increased fungal proportion in high copper (78 versus 65%) and reduced bacterial proportion (35 versus 22 %). The reverse effect was observed during copper chelation, with *S. aureus* proportional CFU increasing to 62%, and *C. albicans* reducing to 38 % of the total biofilm. Therefore, the relative proportions of viable fungi and bacteria within the dual species biofilm is dependent on environmental copper (Figure 4e). To confirm the finding that 50 μm CuSO_4_ reduced *C. albicans* hyphal formation during interaction with *S. aureus*, we imaged biofilms via spinning disk confocal microscopy (Fig 4g-n) and quantified rendered hyphal structures (Fig 4f). The volume of rendered hyphae in dual-species biofilms cultured in 50 μm CuSO_4_ is significantly less than hyphae in dual-species biofilms cultured in standard RPMI (p=0.0002). These findings show that environmental copper alters dual-species biofilm structure and composition by modulating *C. albicans* morphology and the relative proportions of fungi and bacteria

**Figure 4.**
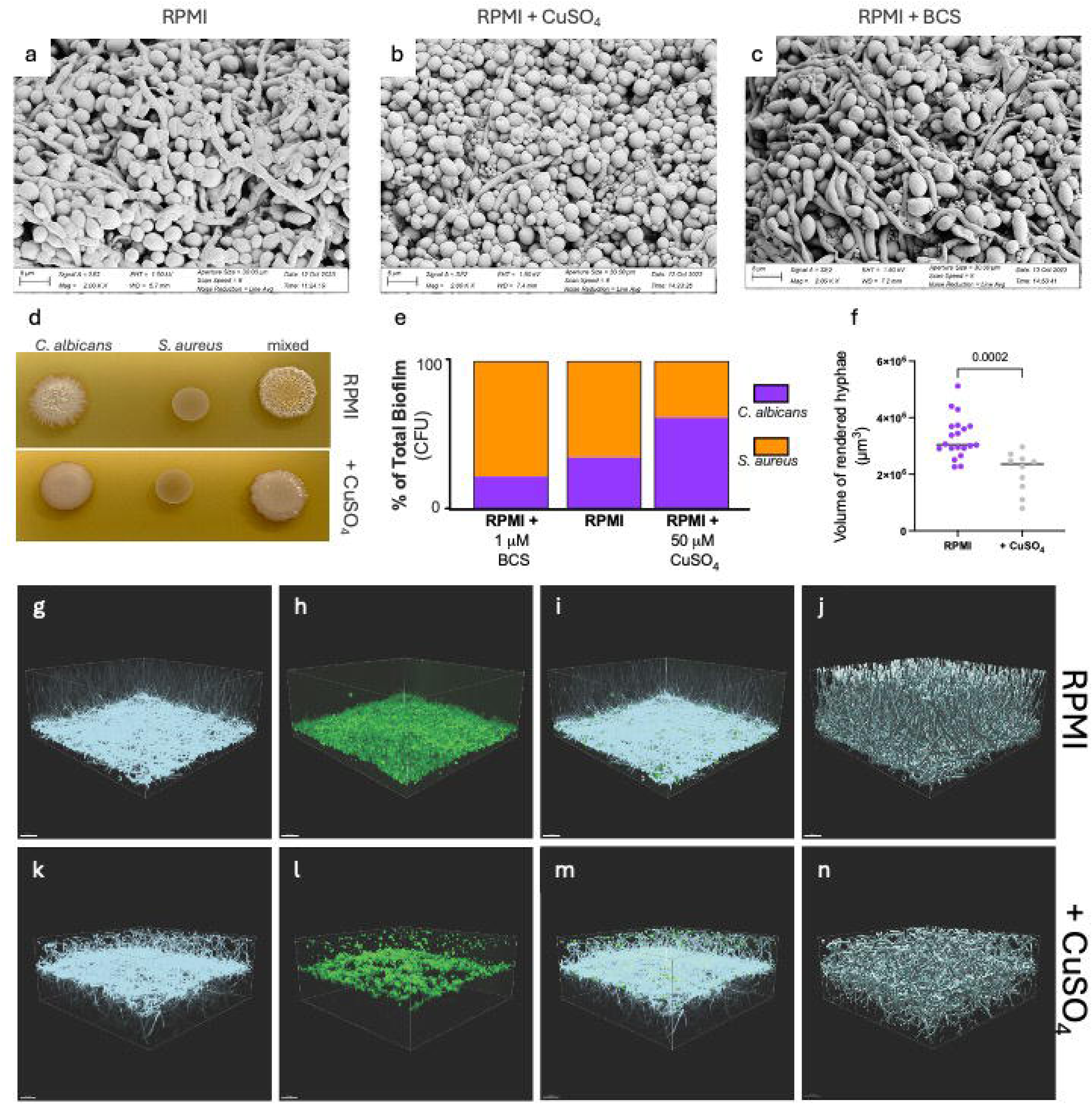
Copper Impacts Dual Species Biofilm Architecture. **a - c** Representative SEM of *C. albicans* single, *S. aureus* single and dual species biofilms in standard (**a**), copper replete (**b**) and copper deplete (**c**) conditions. Biofilms at 200 X magnification (scale bar 5 µM) **d** Representative **i**mages of biofilm cultured on agar pads showing distinct topographical structures of each biofilm, where the top row shows cultures on RPMI and the bottom row shows cultures on RPMI + 50 µM CuSO_4_ **e** Dual species biofilms cultured in the wells of 96 well plates were disrupted, serially diluted and their CFU plated and counted. The proportion of fungal or bacterial cells relative to the total number of cells within the biofilm is shown (n=5) **i** Quantification of fungal volume was performed on biofilms cultured in IBIDI slides and imaged by confocal spinning disk microscopy and analysed IMARIS. Data points represent volume of rendered hyphae (μm^3^) per imaged volume 482 μm x 482 μm x 200 μm. The individual points are individual technical replicates within 3 independent biological experiments (n=3). Statistical significance was determined via a two-tailed student’s *t*-test and a numeric *P value* is shown. **g-n** show representative images of images acquired from spinning disk confocal microscopy. g and k show Calcofluor White labelled fungal cell walls **h** and **l** show GFP signal from *S. aureus.* **i** and **m** show fluorescence overlay of GFP and Calcofluor White channels, and **j** and **n** show corresponding renderings of hyphal volume. The top row shows biofilms in RPMI containing 5% human serum and the bottom row show biofilms in the same media containing 50 µM CuSO_4_.

### Copper Transport via Ctr1 is Necessary for *C. albicans* Single and Dual Species Biofilm

Copper is sensed and transported via distinct mechanisms in fungi and bacteria. To reveal the specific copper machinery contributing to copper use during dual species biofilm growth a panel of *C. albicans* and *S. aureus* copper mutants were subject to biofilm assays (Figure 5). Biofilm assays revealed that bacterial biofilm growth occurs independently of copper transporter CopA (slightly reduced biofilm, *P value* 0.54), and copper related chaperone CopZ (*P value* 0.749) in *S. aureus*. On the fungal side, copper transport was necessary for normal *C. albicans* biofilm formation with biomass reduced to 68% of wild type control biofilms when the *C. albicans* Ctr1 copper transporter was not expressed (crystal violet OD 9.34 +-1.11 versus 6.35 +-6.35; *P value* 0.0241). In experiments crossing the *C. albicans* wild type with the *S. aureus* copper mutants, dual species biofilms were not significantly reduced when CopA or CopZ were absent. When the *C. albicans* copper importer Ctr1 is absent, dual species biofilms fail to reach the levels of even the *C. albicans* Ctr1 single species biofilm. These data demonstrate that fungal copper is required for the synergistic dual species biofilm lifestyle. Given the effect deleting Ctr1 from *C. albicans* has on single and dual species biofilms, we hypothesise fungal copper transport is required for optimal biofilm growth. Furthermore, these data demonstrate that fungal copper transport is integral to the mutualistic interaction of *C. albicans* and *S. aureus* during the biofilm mode of growth.

**Figure 5.**
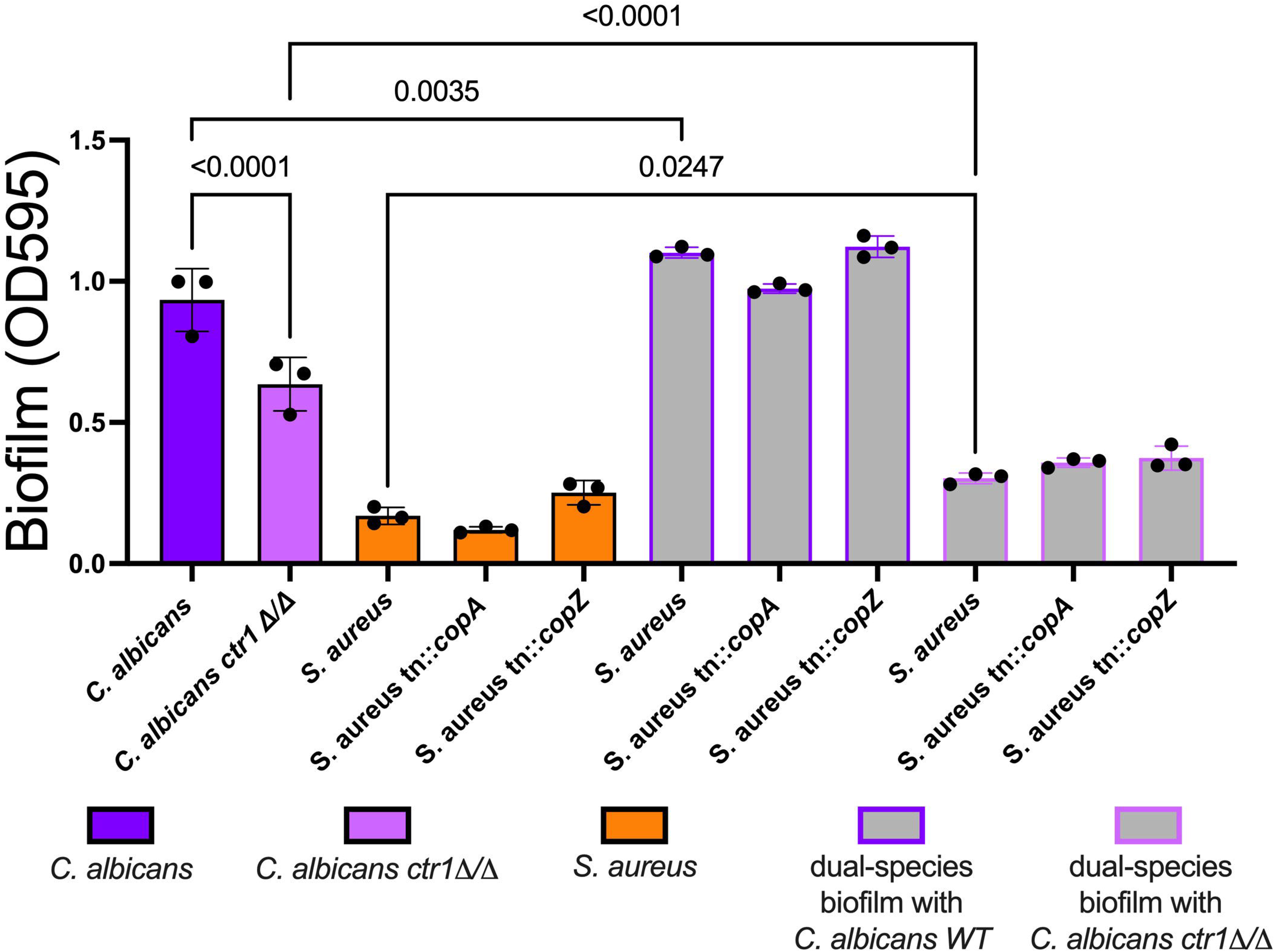
Fungal Copper Transport via Ctr1 is Necessary for *C. albicans* Single and Dual Species Biofilm. Total biomass of single and dual species biofilms was determined via CV assay 48 hours after washing. Dual species biofilms contained *C. albicans* wild type mixed with *S. aureus* wild type or *S. aureus tn::copA* or *S. aureus tn::copZ* (grey bars with purple borders). Additionally, *C. albicans ctr1Δ/Δ* was mixed with *S. aureus* wild type or *S. aureus tn::copA* or *S. aureus tn::copZ* (grey bars with pink borders).Data are shown as absolute absorbance values determined at optical density 595nm where each data point represents the arithmetic mean of a biological replicate, each containing three technical replicates. Error bars represent standard deviation. Statistical significance was determined using a One-Way Anova with Sidak’s multiple comparison tests between all groups. *P* values are shown as numerical values.

### Copper Nanoparticles Disrupt Dual Species Biofilms

Nanoparticles (NPs) represent an emerging therapeutic strategy for combating biofilm-associated infections (45). Despite extensive investigation into the effects of copper nanoparticles on single species biofilms, their impact on polymicrobial communities remains unclear. In light of our findings on the centrality of copper to *C. albicans–S. aureus* dual-species biofilms, we evaluated the efficacy of copper oxide nanoparticles (CuONPs) against these communities. Dual-species biofilms were exposed to a CuONPs at concentrations from 0 to 256 µg /mL, and viability was assessed. CuONPs inhibited biofilm viability in a dose-dependent manner, with an almost 20% reduction observed at 0.5 µg/mL. This inhibitory effect increased until it plateaued at 16 µg/mL, with an approximate 80% reduction in metabolic activity, and no additional reduction was observed up to 256 µg/mL (Figure 6a). This approach demonstrates that CuONP greatly diminish our biofilms, but do not fully eradicate them. To investigate the structural effects of CuONP exposure, scanning electron microscopy was performed on biofilms exposed to the CuONP IC_50_ –2 µg/mL. (Figure 6b-c). While untreated biofilms displayed extensive hyphal scaffolds densely colonized by bacteria (Figure 6b), CuONP-treated biofilms exhibited reduced surface coverage with bacteria, and decreased topographical uniformity (Fig 6c). These alterations mirror the copper-sensitive phenotypes observed in (Fig 4), supporting a model where copper perturbation disrupts fungal-bacterial biofilm integrity.

**Figure 6.**
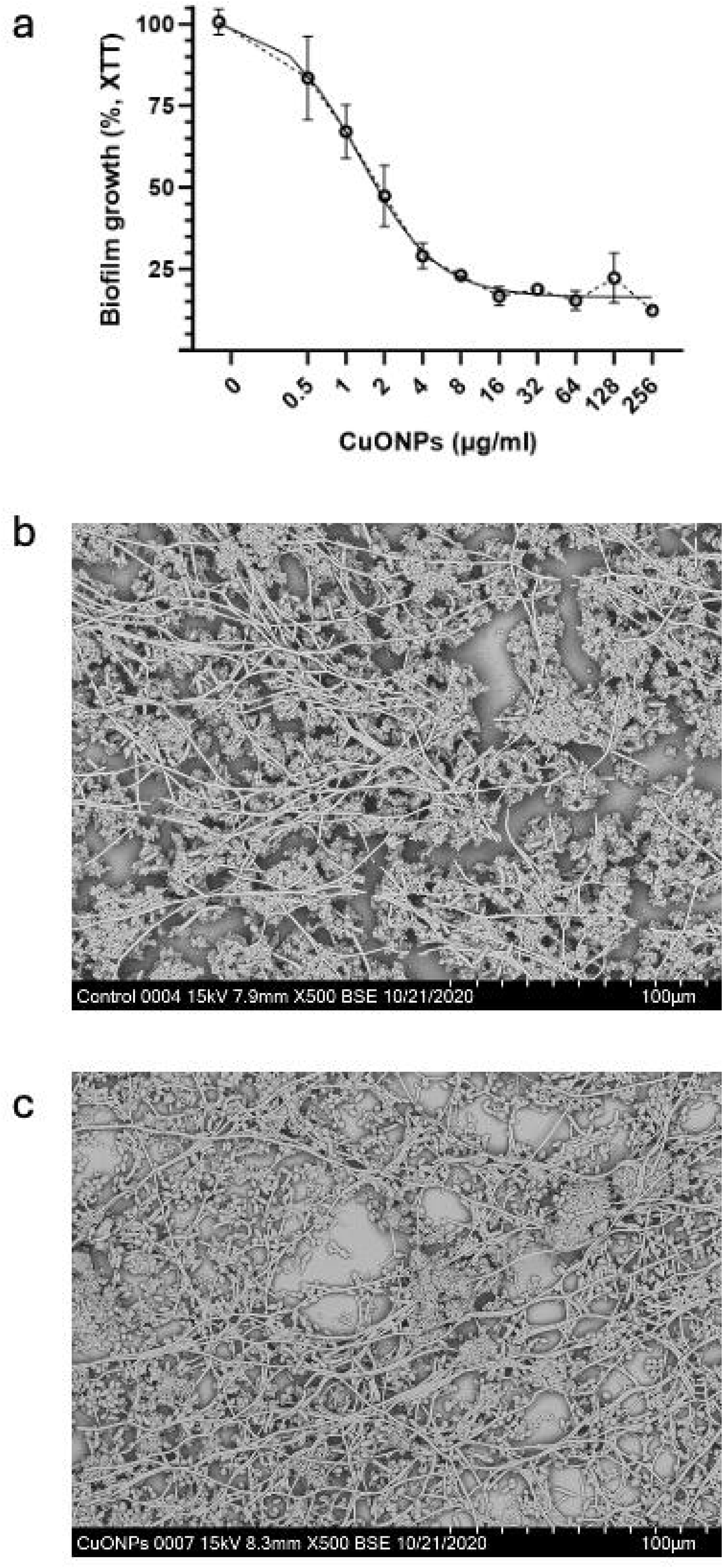
Copper nanoparticles disrupt dual species biofilm. **a** *C. albicans* and *S. aureus* dual species biofilms were exposed to various concentrations of CuONP ranging from 0-256 μg/ml and biofilm metabolic activity was determined via Presto Blue assay. Biofilm metabolic activity is represented as a percentage of untreated biofilm set to 100%. Data shown are the arithmetic mean of three independent biological experiments and error bars represent SEM. The dose-response curve was modelled using a five-parameter logistic (5PL) asymmetric sigmoidal fit in GraphPad Prism, estimating the IC□□ around 2 µg/mL. **b - c** Representative images of dual species biofilm untreated or exposed to CuONP (2 μg/ml) were imaged via SEM.

### A Ctr1 Dependent Propensity of *C. albicans* to Form Mutualistic Biofilm with Bacteria

1. *C. albicans* mixed species biofilms formed with bacteria other than *S. aureus* present a clinical challenge. These biofilms form on medical devices with *Acinetobacter baumannii* (46); in burn wounds with *Staphylococcus epidermidis* (11); and in the lungs wit*h Pseudomonas aeruginosa* (47). Therefore, we sought to determine if there was a widespread impact of copper in the local environment on the ability to form a mutualistic interkingdom biofilm containing *C. albicans*. We employed a panel of Gram-positive *Staphylococci* (*S. aureus MRSA, s. epidermidis*) and Gram-negative (*Escherichia coli, Klebsiella oxytoca, Burkholderia thailandenesis, P. aeruginosa*) bacterial species. Dual species biofilms were formed in standard conditions, then washed and incubated in fresh media. All bacterial species tested (with the exception of *Lactobaccillus crispatus*) formed measurable biofilm in our conditions (Fig 7a). Again, *C. albicans* lacking functional Ctr1 formed poor biofilm compared to its wildtype counterpart (*P value* <0.0001). Dual species biofilms of *C. albicans* and bacteria generally formed mutualistic biofilms of greater biomass than respective single species biofilm (Fig 7b). Dual species biofilms formed with *C. albicans ctr1*Δ*/*Δ and *S. aureus* were again significantly smaller than those formed with *C. albicans* wild type (*P value* =<0.0001). The propensity of *C. albicans ctr1*Δ*/*Δ to form mutualistic dual species biofilm with bacteria was generally less than for wild type (Fig 7c). Interestingly, the dual species biofilm formed of *C. albicans ctr1*Δ*/*Δ and *A. baumannii* was mutualistic, indicating the biofilm gains function in the absence of fungal copper import. These data are summarised in a heatmap (Fig 7d) depicting the propensity of *C. albicans* to form a mutualistic biofilm with bacteria in the presence of functional Ctr1. While wildtype *C. albicans* generally form mutualistic biofilms with bacteria, this is reduced in the absence of Ctr1. Loss of the propensity to form mutualistic biofilm is most significant in the *Staphylococcal* species tested, and *P. aeruginosa*. Therefore, these data point to a prevalent effect of the loss of fungal copper import in *C. albicans* and Gram-positive bacteria interactions. We further extended investigation to a clinical isolate of *C. albicans* and a lab isolate of *Candida parapsilosis* (Supplementary Figure 2) and demonstrated that sub-inhibitory copper effects dual species biofilm formation in these *Candida* species, supporting a broader role for a copper economy in *Candida*–*Staphylococcus* mixed biofilms.

## Discussion

Fungal – bacterial interactions range from antagonistic to synergistic with orchestrated strategies used by microbes to compete, collaborate or survive such as the production of competitive anti-microbial molecules (48) or the modulation of oxidative stress (49). Understanding the molecular basis of interactions between *C. albicans* and *S. aureus* is critical due to their frequent co-isolation in clinical settings and their contribution to a wide spectrum of infections (6,10,13). In this study, we identify copper transport as a previously unrecognised determinant of mutualism between these two pathogens in biofilms. Using proteomics, biofilm assays and microscopy, we demonstrate that copper in the local environment modulates the interaction of *C. albicans* and *S. aureus* during biofilm growth, revealing a copper-dependent axis of mutualism that governs biofilm architecture and biomass. On a proteomic level, we observe a fungal copper-dependent oxidative stress response alongside copper acquisition, while concomitantly bacteria process copper for export from the cell. These data led us to hypothesise that fungal-bacterial community behaviour regarding copper contributed to the mutualistic interaction in biofilm. Our experiments revealed that under both copper replete and deplete conditions, dual species biofilms fail to exceed the biomass of single-species controls, suggesting that a precise balance of copper is required for mutualistic biofilm formation. We observed that copper import to the fungal cell was important for single species biofilm lifestyle, but also dual species biofilm lifestyle. We propose that *C. albicans* responds to the stress of bacterial interaction by increased copper-cofactor superoxide dismutase expression. Our proposed model (Fig 8) puts forth that to achieve community benefit, *S. aureus* contributes to fungal acquisition of the co-factor for Sod1 activity via repression of copper acquisition and shuttling of copper for export via *S. aureus* CopA. This activity could potentially benefit the bacteria by lowering the local environmental copper concentrations which is important as *S. aureus* is sensitive to lower concentration of copper than *C. albicans* (μM versus mM ranges) (50,51). We propose that copper shuttled from bacteria is then taken into the fungal cell via Ctr1 and utilised for Sod1 activity. Additionally, we observe a loss of hyphal structures at elevated copper concentrations and an enrichment of yeast cells (Fig4). As biofilms containing *C. albicans* hyphae provide structure for *C. albicans* and *S. aureus* cell co-localisation at a greater incidence and for longer duration compared to yeast only biofilms (44), we conclude that the loss of mutualism is in part due to reduced co-localisation and interaction. The structural proximity of bacteria adhered to hyphae in our standard conditions, but lost in our copper conditions, likely permits greater interaction leading to development of the mutualistic traits well documented throughout the literature (19,22,52). In support of this model, we quantitatively assessed the impact of copper oxide nanoparticles (CuONPs) on *C. albicans–S. aureus* dual-species biofilms. Scanning electron microscopy revealed that CuONPs disrupted hyphal structures and diminished the uniformity of biofilm architecture, features essential to sustaining mutualism. These findings reinforce the notion that copper modulates not only the biochemical but also the structural basis of fungal-bacterial cooperation and point to CuONPs as promising agents for disrupting polymicrobial biofilms through destabilisation of interkingdom interactions. We extended these observations beyond *C. albicans* and *S. aureus*, to biofilms formed of *C. albicans* with seven additional bacteria and observe a role for fungal copper import in these broad interactions, but particularly in interactions with Gram positives.

**Figure 7.**
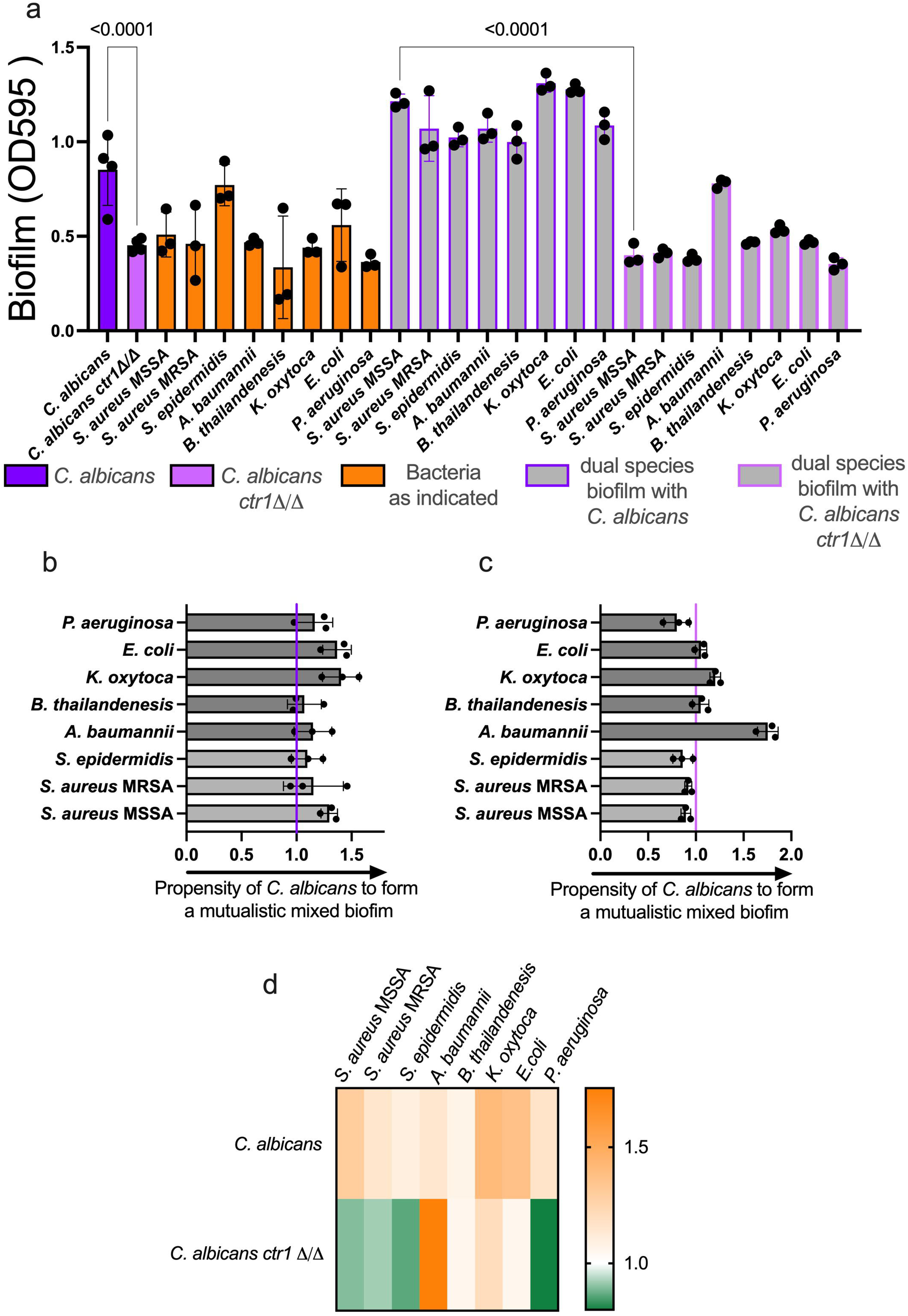
The requirement of fungal copper transport for *C. albicans* – bacterial biofilms. **a** Total biomass of single and dual species biofilms was determined via CV assay after 48 hours of biofilm culture in RPMI. Single species fungal biofilm is shown in purple and single species bacterial biofilms are show in orange. Dual species biofilms are shown in grey, where *C. albicans* wild-type biofilms have a purple border, and *C. albicans ctr1*Δ*/*Δ have a pink border. Data are shown as absolute absorbance values determined at optical density 595nm where each data point represents a biological replicate. Error bars represent standard deviation. Statistical significance was determined using a One-Way Anova with multiple comparisons and Sidak’s correction and *P value*s are shown as numerical values. **b** The propensity of *C. albicans* to form a mutualistic dual species biofilm with various bacteria was determined by comparing the total biomass of single species fungal biofilm to the biomass of dual species biofilms to generate a value representing the propensity to form mutualistic dual species biofilms with bacteria (OD595 of dual species biofilm / OD595 single species biofilm = propensity to form a mutualistic biofilm) whereby a value of 1 (indicated by a purple line) indicates a biofilm the same as the single species biofilm, a value less than 1 indicates an antagonistic biofilm and a value greater than 1 indicates a mutualistic biofilm. This calculation was performed using the data from (a). Dual species biofilm formed of Staphylococcal species is shown in light grey, with non-staphylococcal bacterial shown in dark grey. **c** The propensity of *C. albicans ctr1*Δ*/*Δ to form a mutualistic dual species biofilm with various bacteria was determined as in (b). Here the value of 1 is marked with a pink line. This calculation was performed using the data from (a). Dual species biofilm formed of Staphylococcal species is shown in light grey, with non-staphylococcal bacterial shown in dark grey **d** A heatmap showing the relative propensity of *C. albicans* wild-type (top row) or *C. albicans ctr1*Δ*/*Δ (bottom row) to form a mutualistic dual species biofilm with bacteria is shown.

**Figure 8.**
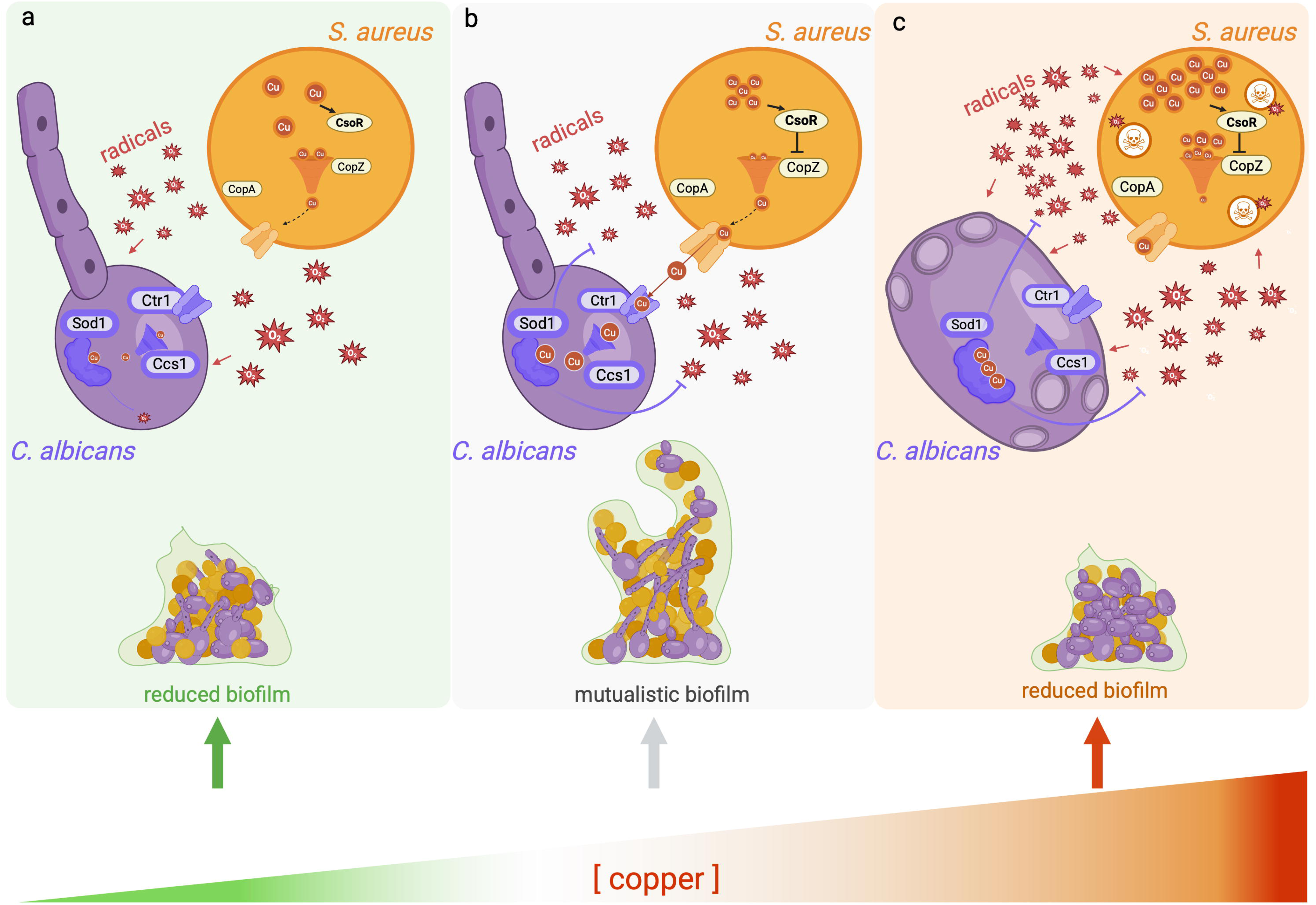
The Copper Economy: A schematic representation of *C. albicans* - *S. aureus* dual species biofilm formation across gradients of copper. (a) At low concentrations of copper, achieved using copper chelation, *C. albicans* and *S. aureus* form biofilms of similar biomass to their single species counterparts. At low copper, dual species biofilms are enriched for *S. aureus* cells. (b) At optimal conditions for mutualistic growth, biofilms are of greater biomass and metabolic activity than single species counterparts. Our model proposes that *S. aureus* exports copper which is then acquired by *C. albicans* via Ctr1 to use as a co-factor for copper-specific detoxification of the free radicals generated by the fungal-bacterial interaction. This mutually beneficial use of copper within the biofilms permits mutualistic biofilms. (c) However, at high concentrations of copper achieved by addition of CuSO_4_, dual species biofilms form similar biomass as single species counterparts and bacterial cells are diminished. Our model proposes that elevated concentrations of copper are toxic to *S. aureus* which cannot export copper to *C. albicans* due to increased intrinsic oxidative stress, cell damage and cell death. Finally, we propose that a copper economy within *C. albicans* and *S. aureus* biofilms contributes to mutualistic interactions.

Copper is both essential and toxic to fungi, bacteria and humans. It is a redox-active metal ion, vital to numerous cellular activities but toxic in high quantities. As a result, copper exists in either a reduced state Cu(I), or an oxidized state Cu(II), allowing it to interact with a wide variety of biological ligands where it has fundamental structural and catalytic roles for example Cu/Zn superoxide dismutase (SOD), cytochrome oxidase and methane mono-oxidase (53). During infection, the use of copper by pathogens and the human host is dynamic depending on the specific niche, environmental conditions and physiological signals. For example, copper is a cofactor for fungal Sod1, and microbes isolated from patients have been shown to switch between use of copper or manganese co-factor SODs (Sod1 or Sod3, respectively) depending on availability of cofactors within the human host (54,55). During *C. albicans* infection, mice kidneys initially increase copper concentration up to 24 hours post infection, followed by prolonged depletion, while in the blood there is an initial increase which is sustained throughout a 96-hour infection. Simultaneously, professional phagocytes – the cells that engulf foreign particles– build mitochondrial copper stores to become activated (56) and concomitantly weaponise copper stores to expose invading pathogens to toxic copper concentrations (57). Contact with copper, such as copper surfaces, damages the cell envelope (58) and excess intracellular copper leads to generation of hydroxy radicals which damage nucleic acids and proteins (59). Thus, in response to host-imposed copper, fungi and bacteria have developed stringent transport mechanisms and coordinate reduction in expression of copper-requiring proteins (60,61).

In line with a classic excess-copper response described for *S. aureus* (50), we detected elevated Clp proteases and thioredoxin-mediated detoxification. Unexpectedly, however, rather than down-regulating protein synthesis—as Baker and colleagues observed in planktonic cells—we observed a marked increase in total *S. aureus* protein content and enrichment of ribosomal proteins within dual species biofilms. Baker *et al*. also reported copper-mediated biofilm inhibition via suppressed Agr and Sea activity, yet under our conditions single-species *S. aureus* biofilms were unaffected. We did, however, detect Rot modulation, which—through repression of Agr—could influence biofilm development. We attribute these discrepancies to differences in model system (planktonic versus mature biofilm) and timelines. Collectively, these findings reveal that copper-responsive pathways not only safeguard *S. aureus* against copper toxicity but also facilitate its mutualistic coexistence with fungi in mixed-species biofilms.

We observed that copper-based mutualism is most pronounced when *C. albicans* interacts with *Staphylococci*, compared to Gram-negative bacteria. The periplasms of Gram-negative bacteria pose an additional barrier to copper export and detoxification, and as a result, these bacteria have developed additional systems to deal with high copper concentrations (62). We speculate that copper plays less of a role in the interaction of *C. albicans* with Gram negative pathogens, as the community use of copper has developed differently in these organisms due to the burden of bacterial copper export and additional periplasmic copper detoxification.

Fungal Sod1 is the most differentially expressed protein by *C. albicans* in dual species biofilms with *S. aureus*, indicating its important role in the interaction between these microbes in the biofilm mode of growth. Specifically, Sod1is a super oxide dismutase which requires copper as a catalytic co-factor. It is located between the cytosol and the mitochondrial intermembrane space and relay between these two cellular compartments requires the Sod1 copper chaperone, CCS (63), which is also highly expressed in mixed biofilms and functions to incorporate copper into Sod1 (64). Sod1 functions to scavenge oxygen radicals by disproportionation of superoxide anion free radicals into less toxic hydrogen peroxide, and molecular oxygen (65). Generally, Sod1 expression is induced in response to general or oxidative stress often caused by aerobic respiration or reactive oxygen species (ROS) generation. Increased Sod1 expression may also contribute to the azole tolerance phenotypes of mixed biofilms by offsetting the toxicity of ROS generated during exposure to drugs (66).

*C. albicans* Als3 is a multifunctional protein expressed on the surface of hyphae and implicated in biofilm formation (67), attachment of *S. aureus* to hyphal structures (19) and absorption of copper (68). Despite the connection of this protein’s function with many aspects of our study, we did not detect differential expression of Als3 with our proteomic approach, potentially as it is regulated at the transcriptional level. However, work exploring if our observations are Als3 related is underway.

*C. albicans* and *S. aureus* collaborate on the use of copper for community benefit. Whether coordinated copper use is an active process to create a non-toxic copper niche for the bacteria, or if Sod1 expression occurs as a result of excess copper remains to be determined. Expression of *S. aureus* copper export and chaperone activity suggest *S. aureus* responds to a copper-rich environment when in biofilm co-culture with *C. albicans*. Indeed, these two organisms have distinct tolerances for copper with *C. albicans* tolerating copper at concentrations 200-fold higher than *S. aureus* (50,51). However, single species biofilms did not respond to extremes of environmental copper in the same way as dual species biofilms did, and we can’t foresee any mechanism through which copper levels could become elevated within the biofilms. Furthermore, as our data were captured during mutualistic growth, we are working on the hypothesis that the microbes are engaging in activity they mutually benefit from. Copper quantification via ICP-MS, temporal transcriptomic and reporter-based approaches are a current focus of our lab and could clarify the dynamics of copper handling between these organisms.

Baseline concentrations of copper vary within the human host depending on anatomical niche, genetic predispositions and infection progression (54,69,70). Furthermore, exposing invading pathogens to high concentrations of copper within the phagolysosome is a strategy employed by immune cells to kill microbes (55,57). Therefore, commensal or pathogenic microbes will be exposed to gradients of copper throughout their natural lifestyles. Our observation that hyphal morphology sustains dual species biofilm formation under standard conditions but is lost in copper-replete environments supports a model in which *C. albicans –S. aureus* interactions are context-dependent—coexisting silently in the microbiome and becoming cooperative or mutualistic only under virulence-inducing conditions where copper levels are elevated. It is intriguing to speculate that copper may serve as a molecular signal influencing this transition. Importantly, the relevance of copper to infection biology extends beyond *in vitro* observations; Elevated copper levels have been observed in cystic fibrosis (CF) patients (69), a population at high risk for polymicrobial infections (71). Whether altered copper homeostasis contributes to increased co-infection susceptibility remains an open question. If copper dysregulation promotes mutualistic interactions between pathogens, host copper status could represent an unappreciated risk factor for poor co-infection outcomes.

Taken together, our findings position copper as an integral element in the mutualistic relationship between *C. albicans* and *S. aureus*. By revealing micronutrient coordination between *C. albicans* and *S. aureus* that supports biofilm growth, this work highlights copper as a promising target for disrupting interkingdom cooperation. By probing copper homeostasis, signalling, and stress adaptation in polymicrobial biofilms, we may uncover new therapeutic avenues to combat recalcitrant co-infections. Indeed, copper-alloy surfaces in intensive care units (ICU) have been linked with up to 83% reduction in microbial burden (72) and copper-based therapies which exploit Sod1 and Ctr1 activity are already in clinical use (73). Within the context of well-established use of copper drugs and antimicrobial coatings, our work opens the door to more informed applications of metal-based therapies to address problems caused by biofilms across healthcare, agriculture, and industry.

Finally, development of mutualism in genetically unrelated populations is thought to begin with the trading of byproducts (74). Here, we provide evidence for the trade of copper in a fungal-bacterial population. We posit that, within our system, copper is a product of lower value to bacteria than it is to fungi, and so it is exchanged for the net benefit of greater detoxification. Resulting from this exchange, both trade partners profit from a larger, more resilient biofilm. We demonstrate that mutualism in our system functions along an axis of copper availability. Thus, our work depicts a landscape within *C. albicans* – *S. aureus* biofilms where copper is a valuable commodity utilised for the good of the community. We have termed this concept the “Copper Economy”.

## Materials and Methods

### Media and Growth Conditions

*Candida* strains were routinely cultured in Yeast Peptone Dextrose (YPD – Sigma-Aldrich) supplemented with 2% glucose (Sigma-Aldrich). *Staphylococcus* strains were routinely cultured in Tryptic Soy Broth (TSB). Single colonies were transferred from agar plates to a 5 ml broth in a 50 mL falcon and cultured for 20-24 hours with shaking at 180 rpm. *C albicans* was cultured at 30°c, while *S. aureus* was cultured at 37°C. Prior to experiments, stationary cultures were washed three times in 1 X PBS and resuspended in RPMI-1640 with Glutamax (Fisher Scientific) with 5% human serum (Cambridge Biosciences), which had been heat inactivated for 30 minutes at 55°C. *C. albicans* was selected for on YPD agar containing 2% glucose and 50 μg / ml vancomycin (Sigma-Aldrich), while *S. aureus* was selected for on TSA containing 10 μg / ml fluconazole (Sigma-Aldrich). Gram-negative bacteria were cultured in LB media (Sigma-Aldrich). Strains cultured in this study are listed in Table 2

**Table 2.** Strains used in this study (54,75–80)

| Name | Description | Reference |
| --- | --- | --- |
| *C. albicans* SC5314 | lab reference stain, wild type, blood isolate | Gillum *et al* 1984 |
| *C. albicans* *ctr1* $\Delta/\Delta$ | *C. albicans* SC5314 *ctr1* deletion strain | Mackie *et al* 2016 |
| *C. albicans* SCD0070 | respiratory isolate | This study |
| *C. parapsilosis* | lab reference stain, wild type | ATCC22019 |
| *S. aureus* SH1000 | lab reference stain, wild type | Horsburgh *et al* 2002 |
| *S. aureus* GFP | tetracycline resitant GFP expressing *S. aureus* | Boldock *et al* 2018 |
| NE590 | JE2 harbouring a transposon at *copA*; *JE2 tn::copA* | Fey *et al* 2013 |
| NE1660 | JE2 harbouring a transposon at *copZ*; *JE2 tn::copZ* | Fey *et al* 2013 |
| *S. aureus* SH1000 *tn::copA* | SH1000 phage transduced to harbour the NE590 transposon, | This study |
| *S. aureus* SH1000 *tn::copZ* | SH1000 phage transduced to harbour the NE1660 transposon, | This study |
| *S. aureus* JE2 | lab reference stain, wild type, MRSA | Fey *et al* 2013 |
| *S. epidermidis* | lab reference stain, wild type | ATCC 35984 |
| *A. baumannii* | lab reference stain, wild type | ATCC19606 |
| *B. thailandensis* | lab reference stain, wild type | Kraus *et al* 2024 |
| *K. oxytoca* | voice prosthesis isolate | This study |
| *E. coli* | BW25113, lab reference strain, wild type | Datsenko *et al* 2000 |
| *P. aeruginosa* | lab reference stain, wild type | ATCC10145 |

### *S. aureus* genetic manipulation

*S. aureus* JE2 Nebraska Transposon Mutant Library (NTML) ((78)) strains were crossed into our lab SH1000 strain via lambda phage transduction. Transposon mutants are listed in Table 2. Bacteriophage □11 were propagated on the indicated mutants as described elsewhere ((81)). Briefly, 100 μL of □11 (∼1 × 10^10^ bacteriophage), and 100 μL of stationary phase NTML strain overnight donor strain were combined by inversion and incubated statically at room temperature for ∼15 hours. Resulting bacteriophage was passed through a 0.45-μm filter syringe and stored at 4°C until use. Transduction was performed as described elsewhere ((24)). Briefly, a stationary *S. aureus* SH1000 culture was diluted 1:100 in TSB and incubated at 37°C for 1 hour. Cells were pelleted and resuspended in 0.5 mL TSB, followed by the addition of 40 μL CaCl_2_ (10 mg/mL) and 100 μL of phage (from the step above). Cells with bacteriophage were incubated for 10 minutes at room temperature and 30°C for 35 minutes. Cells were pelleted, resuspended in 5 mL TSB and incubated 37°C for 90 minutes, with shaking. Multiple 100 μL volumes were plated on TSA containing erythromycin (5 ug/mL) to select for mutant colonies.

### Biofilm culture

Cells from overnight cultures were harvested by centrifugation, washed three times in 1 x PBS and resuspended in RPMI-1640 with Glutamax (Fisher Scientific). Cells were counted using a ViCell Blu (*Candida*) or haemocytometer (*Staphylococcus*) and normalised to 2 x 10^7^ cells/ mL in RPMI-1640 containing 5% human serum (HA). A 50 μL volume of each species was transferred to the well of a 96 well plate, so that single species biofilms contained 50 μL of cell suspension and dual species biofilms contained 100 μL of cell suspension. Media was added to each well to achieve a final volume of 200 μL. Media without cells acted as a biofilm negative control. Plates were incubated statically at 37°C with 5% CO_2_ for 4 hours. After this time, non-adherent cells were removed by careful washing of the wells with sterile 150 μL volumes of 1 X PBS, three times. Adherent biofilms were replenished with 200 μL RPMI-1640 with 5% human serum and incubated at 37°C with 5% CO_2_ for the indicated time points. Where indicated, the media to replenish adherent biofilms contained bathocuproinedisulfonic acid disodium salt (BCS, Sigma-Aldrich), copper sulphate (CuSO_4_, Sigma-Aldrich), Magnesium chloride (Sigma-Aldrich) or Zinc Chloride (Sigma Aldrich).

### Biofilm quantification

Following incubation, media and non-adherent cells were removed by pipetting and washing with 150 μL volumes of PBS three times. Biofilms were dried for one hour at room temperature, then stained with 200 μL 0.1% (w/v) Crystal Violet solution prepared in dH_2_0. The stain was allowed penetrate the biofilm at room temperature for 30 minutes, then was removed and washed away with three 150 μL volumes of PSB. Stained biofilms were dissolved in 200 μL 10% acetic acid (v/v) prepared in dH_2_0. Suspensions were transferred to fresh 96 well plates for measurement, and when required, were diluted in dH_2_0. Crystal violet, as a measure of total biofilm biomass, was measured at OD595nm.

### Biofilm metabolic activity

Biofilms were cultured in 96 well plates and incubated as described. At the indicated time points, biofilms were subject to the XTT Cell Viability kit (Sigma-Aldrich), or the Presto Blue kit (Sigma-Aldrich) without deviation from manufacturer’s instructions. Presto blue assay fluorescence was measured after 15 minutes incubation at 37°C in a Synergy II Spectrophotometer (Ex 530nm/Em 590nm).

### Copper nanoparticle antibiofilm assay

To determine the effect of CuONPs on mixed biofilm, Copper Oxide Nanoparticles (<100 nm) were acquired from SkySpring Nanomaterials (USA) and incubated with biofilms. Microbial cultures were grown overnight in TSB and YPD, respectively. Dual-species biofilms were formed in a 1:1 RMPI/biofilm broth mixture, seeded at 10C cells/mL for *C. albicans* and 10C cells/mL for *S. aureus*, and incubated in 96-well plates with CuONPs in a 0.5–256 µg/mL concentration gradient. After 24 h at 37 °C, biofilms biofilm metabolic activity was recorded.

### Biofilm protein extraction

For proteomics, biofilms were cultured as described but in 24 wells plate to allow for efficient biofilm disruption and efficient protein extraction. At 48 hours, biofilms were agitated using a yellow 200 μL pipette tip and transferred to 1.5 mL polypropylene tubes, centrifuged to pellet cells at 3,000 g for 10 minutes, then resuspended in 200 μL Yeast Buster Protein Extraction Reagent (Millipore) containing 200 μg/mL lysostaphin and incubated for 30 minutes at 37°C with gentle agitation. Insoluble debris was removed by centrifugation at 3,000 g for 5 minutes. Protein concentration was determined using the BCA assay kit (Millipore Merck). Equal volumes (50 μL) of protein were analysed.

### TMT labelling and high-pH reverse-phase chromatography

Samples were subjected to FASP-based proteolytic digestion, essentially as described in (82), using 30kDa molecular weight cut-off filters (VWR International Ltd. UK) and digestion with trypsin (1 μg trypsin 1; 37°C, overnight). Post-digestion, peptides were desalted using C18 tips, according to the manufacturer’s instructions (Thermo Fisher Scientific, Loughborough, LE11 5RG, UK), evaporated to dryness and resuspended in 100mM TEAB. Samples were then labelled with Tandem Mass Tag (TMT) ten plex reagents according to the manufacturer’s protocol (Thermo Fisher Scientific) and the labelled samples pooled and desalted using a SepPak cartridge, according to the manufacturer’s instructions (Waters, Milford, Massachusetts, USA). Eluate from the SepPak cartridge was evaporated to dryness and resuspended in buffer A (20 mM ammonium hydroxide, pH 10) prior to fractionation by high pH reversed-phase chromatography using an Ultimate 3000 liquid chromatography system (Thermo Fisher Scientific). In brief, the sample was loaded onto an XBridge BEH C18 Column (130Å, 3.5 µm, 2.1 mm X 150 mm, Waters, UK) in buffer A and peptides eluted with an increasing gradient of buffer B (20 mM Ammonium Hydroxide in acetonitrile, pH 10) from 0-95% over 60 minutes. The resulting fractions (concatenated into 8 in total) were evaporated to dryness and resuspended in 1% formic acid prior to analysis by nano-LC MSMS using an Orbitrap Fusion Lumos mass spectrometer (Thermo Scientific).

### Nano-LC mass spectrometry

High pH RP fractions were further fractionated using an Ultimate 3000 nano-LC system in line with an Orbitrap Fusion Lumos mass spectrometer (Thermo Scientific). In brief, peptides in 1% (vol/vol) formic acid were injected onto an Acclaim PepMap C18 nano-trap column (Thermo Scientific). After washing with 0.5% (vol/vol) acetonitrile 0.1% (vol/vol) formic acid peptides were resolved on a 250 mm × 75 μm Acclaim PepMap C18 reverse phase analytical column (Thermo Scientific) over a 150 min organic gradient, using 7 gradient segments (1-6% solvent B over 1min., 6-15% B over 58min., 15-32%B over 58min., 32-40%B over 5min., 40-90%B over 1min., held at 90%B for 6min and then reduced to 1%B over 1min.) with a flow rate of 300 nl min^−1^. Solvent A was 0.1% formic acid, and Solvent B was aqueous 80% acetonitrile in 0.1% formic acid. Peptides were ionized by nano-electrospray ionization at 2.0kV using a stainless-steel emitter with an internal diameter of 30 μm (Thermo Scientific) and a capillary temperature of 300°C. All spectra were acquired using an Orbitrap Fusion Lumos mass spectrometer controlled by Xcalibur 3.0 software (Thermo Scientific) and operated in data-dependent acquisition mode using an SPS-MS3 workflow. FTMS1 spectra were collected at a resolution of 120 000, with an automatic gain control (AGC) target of 200 000 and a max injection time of 50ms. Precursors were filtered with an intensity threshold of 5000, according to charge state (to include charge states 2-7) and with monoisotopic peak determination set to Peptide. Previously interrogated precursors were excluded using a dynamic window (60s +/-10ppm). The MS2 precursors were isolated with a quadrupole isolation window of 0.7m/z. ITMS2 spectra were collected with an AGC target of 10 000, max injection time of 70ms and CID collision energy of 35%. For FTMS3 analysis, the Orbitrap was operated at 50 000 resolution with an AGC target of 50 000 and a max injection time of 105ms. Precursors were fragmented by high energy collision dissociation (HCD) at a normalised collision energy of 60% to ensure maximal TMT reporter ion yield. Synchronous Precursor Selection (SPS) was enabled to include up to 10 MS2 fragment ions in the FTMS3 scan.

### Proteomics data analysis

The raw data files were processed and quantified using Proteome Discoverer software v2.4 (Thermo Scientific) and searched against the UniProt *Staphylococcus aureus* [93061] database (downloaded December 2022; 2889 sequences) and the Uniprot *C. albicans* [237561] database (downloaded December 2022; 6031 sequences) using the SEQUEST HT algorithm. Peptide precursor mass tolerance was set at 10Cppm, and MS/MS tolerance was set at 0.6CDa. Search criteria included oxidation of methionine (+15.995Da), acetylation of the protein N-terminus (+42.011Da) and methionine loss plus acetylation of the protein N-terminus (-89.03Da)as a variable modifications and carbamidomethylation of cysteine (+57.021Da) and addition of the TMT mass tag (+C229.163Da) to peptide N termini and lysine as fixed modifications. Searches were performed with full tryptic digestion, and a maximum of two missed cleavages were allowed. The reverse database search option was enabled, and all peptide data were filtered to satisfy a false discovery rate of 5%. <u>Normalised unpaired analysis:</u> Protein abundance data were normalised by species; the sum abundance of all proteins of a single species was calculated relative to all proteins of that same species. A normalisation sample was applied to each factor which brought the abundance totals in line with the maximum total. For example, *S. aureus* normalisation was not applied to *C. albicans* and vice versa, but was applied to *S. aureus* single species biofilm and also to *S. aureus* proteins in the dual species biofilm.

### Scanning electron microscopy

For scanning electron microscopy (SEM), biofilms were prepared by spotting 20 μL of fungal, bacterial, or fungal-bacterial mixed cultures on RPMI-1640 agar (Thermo Fisher Scientific) containing 5% human serum and CuSO_4_or BCS as indicated. Plates were incubated for 48 hours at 37°C and 5% CO_2_. Biofilm agar pads were excised using a sterile scalpel and transferred to single well of a 12 well plate. Biofilms were fixed for 60 minutes by immersion in 2% glutaraldehyde, 2% paraformaldehyde in 0.1M sodium cacodylate buffer (pH 7.2). Following three 5-minute washes in buffer, samples were post-fixed in 1% aqueous osmium tetroxide for 60 minutes, then washed three times for 5 minutes in deionized water before dehydration in a graded ethanol series (30%, 50%, 70%, 80%, 90%, 95% for 10 min each, and 100% ethanol for 2 x 15 min). Samples were then incubated in 100% hexamethyldisilazane (HMDS) for 3 minutes before air drying. Dried samples were carefully mounted on aluminium stubs with the help of adhesive copper tabs, then sputter coated with 10 nm gold/palladium. Biofilms were imaged using a Zeiss GeminiSEM 500 operated at 1.5kV.

### Confocal microscopy and IMARIS analysis

For confocal microscopy, biofilms were cultured as described, in IBIDI 8-chamber slides and wild type *C. albicans* and *S. aureus* GFP were used. At the indicated timepoint, media was removed by gentle pipetting and biofilms were washed three times in 1 X PBS. Biofilms were fixed for 30 minutes in 4% paraformaldehyde in 1 X PBS, at room temperature. Fixation solution was removed, biofilms washed once, then subject to 10 minutes 1mg/mL calcofluor white (CFW) in 1 X PBS. Staining was removed, biofilms were washed and stored in 1 x PBS at 4°C. Refractive index adjustment was achieved with 30 min incubation in 38% iohexol (Sigma, UK) in TRIS-EDTA buffer pH 7.4. Biofilm samples were imaged using a confocal spinning disc microscope Dragonfly 505 (Andor, Oxford Instruments, UK). Z-stack was achieved with 1 μm between optical sections and 200 μm height was acquired. Rendering of hyphal volume was performed using Imaris software.

### Statistics

Experiments were performed independently at least three times. The exact number of experiments are provided by individual data points in the figures, and in the figure legends. Statistical analyses were performed using Prism GraphPad 10 and the specific tests are specified in the figure legends.

## Acknowledgements

We thank Alexandra Brand, Cameron Bedford, Alistair Brown, Ian Leaves, Ben Temperton, Stefano Pagliara, Maisem Laabei, Simon Foster, and Mariana Tinajero-Trejo for generously providing strains that either feature in this manuscript or contributed to its development. We are grateful to Paulina Cherek for assistance with electron microscopy sample preparation, and to the Mycoscopy Lab at the MRC CMM, led by Darren Thomson. RVB thanks Jose L. Lopez-Ribot at the University of Texas at San Antonio for access to facilities and instrumentation. Special thanks to Adilia Warris and Elaine Bignell for their valuable discussions throughout the course of this work and helpful comments on the manuscript. We are grateful to Elaine Bignell for coining the term “copper economy.”

