## Supplemental figures 1 and 2 for "Copper Driven Mutualism of *Candida albicans* and *Staphylococcus aureus* Interkingdom Biofilms"

**Supplemental Data**

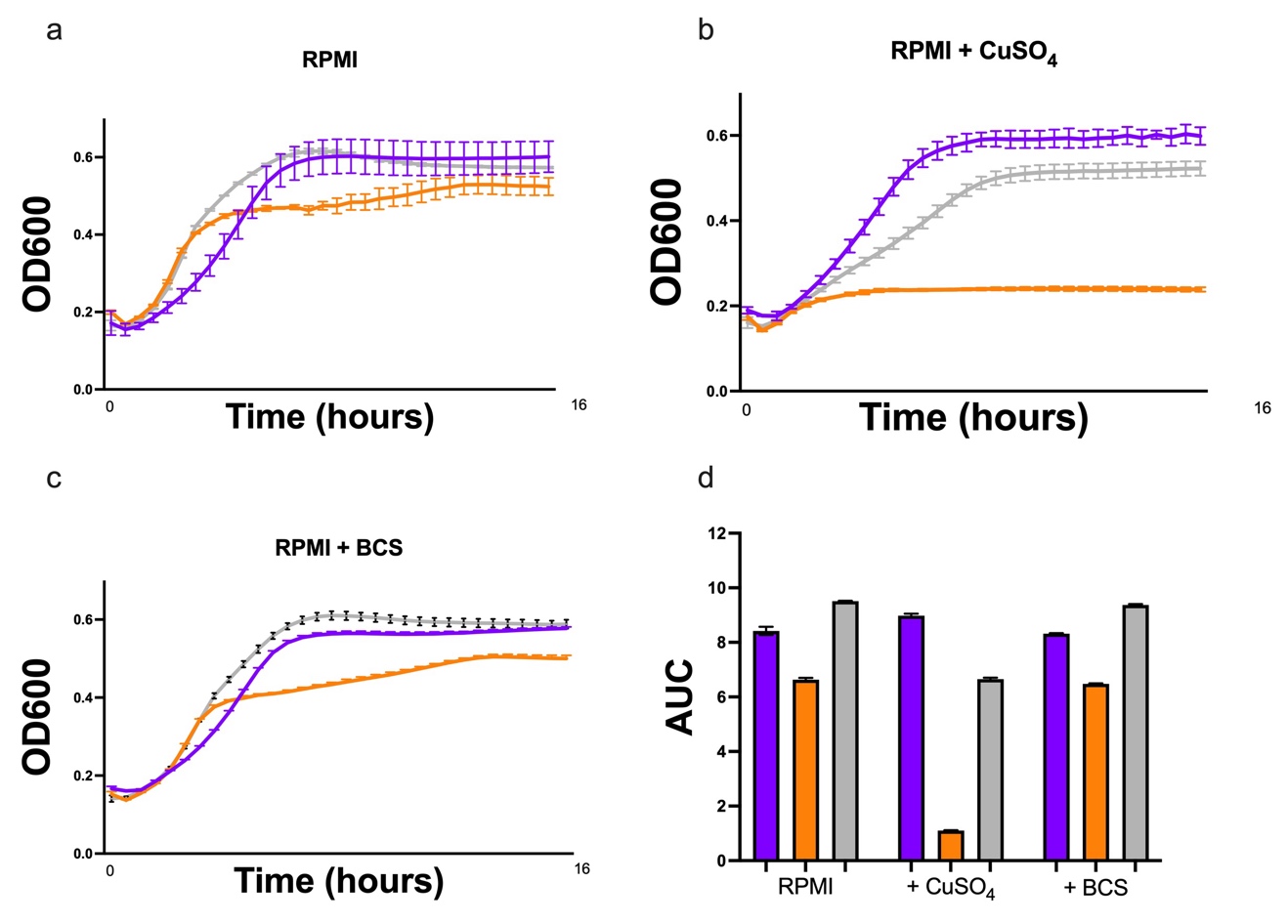

**Figure S1 Single and dual species planktonic culture in RPMI and copper conditions** | **a** OD600 of planktonic cultures in RPMI where *C. albicans* is shown as a purple line, *S. aureus* shown as an orange line and co-cultures of both shown as a grey line. Overnight cultures we washed, diluted 1:1000 in RPMI and transferred in 200 μL to 96 well plates were OD600, indicative of microbial growth, was recorded at 30-minute intervals up to 16 hours on a Tecan M-Plex plate reader. Growth curves were also performed in **b** 50 µM CuSO_4_ and **c** 1 µM BCS. n=3. **d** Area under the curve (AUC) was calculated for each culture in **a**, **b** and **c**. Bars represent arithmetic mean AUC for *C. albicans* (purple), *S. aureus* (orange) and co-cultures (grey).

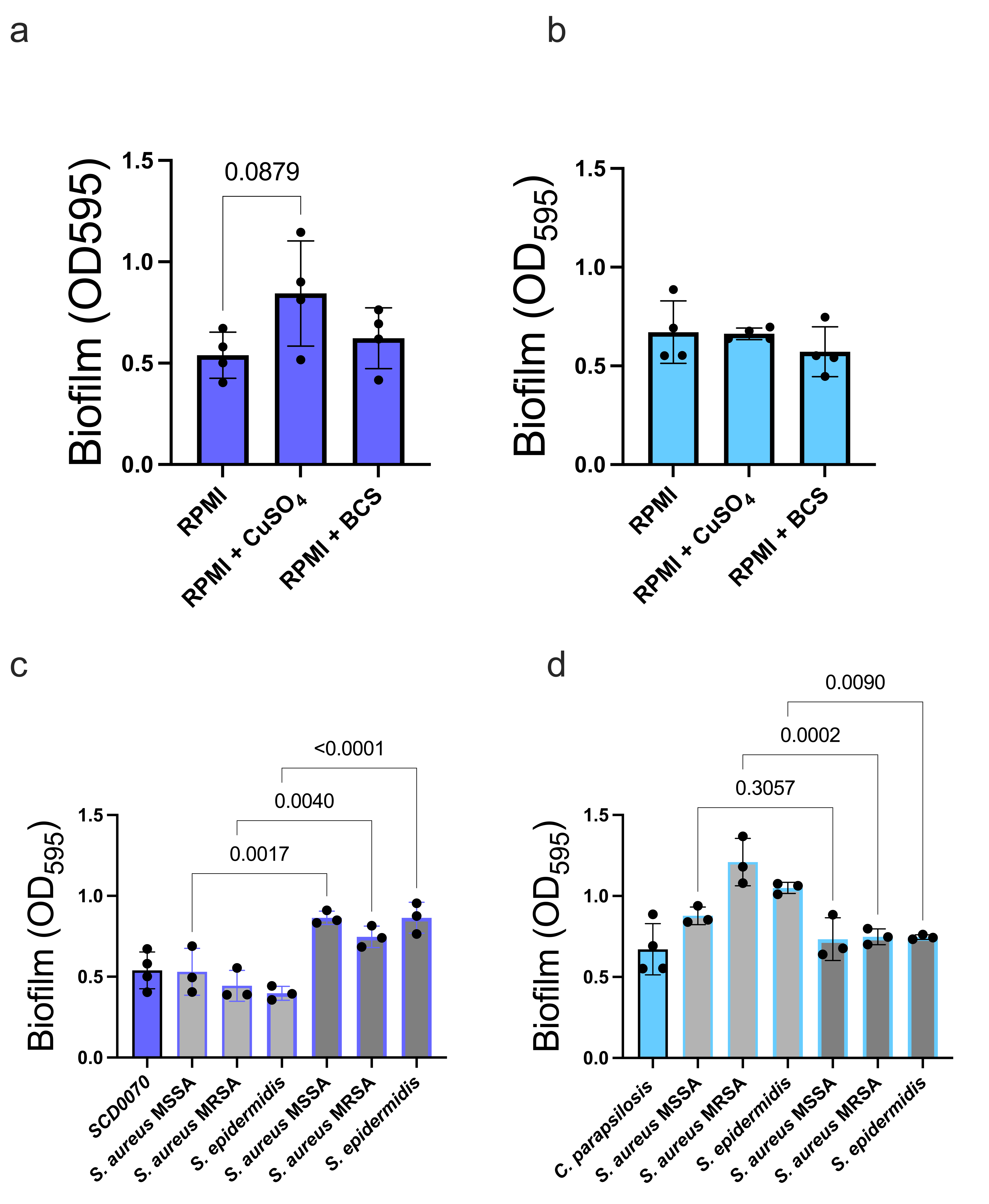

**Figure S2 A Copper Economy of *Candida* – *Staphylococci* Species Biofilms | a and b** To investigate if copper contributes to the ability of non-SC5314 *C. albicans* species and non-albicans species, we quantified biofilm in a *C. albicans* clinical isolate which we’ve named SCD0070 and *Candida parapsilosis*. While there is no significant effect of CuSO_4_ or BCS at the concentrations used throughout this work on either strain, there is a trend for 50 µM CuSO_4_ to increase single species biofilm formation in *C. albicans* SCD0070 (*P value* 0.0879). **c and d** The formation of dual species biofilm with Staphylococci species *S. aureus* SH1000 (MSSA), *S. aureus* JE2 (MRSA) and *S. epidermidis* was quantified in RPMI (light grey bars) and in RPMI with 50 µM CuSO_4_ (dark grey bars) There was no change in biofilm formation when C. albicans SCD0070 was mixed with the staphylococci species in RPMI. However, there were significant increases in biofilm when these species were mixed in the presence of 50 µM CuSO_4_. Conversely, *C. parapsilosis* formed mutualistic biofilm with staphylococci species in RPMI and this was abrogated in the presence of copper, mirroring the observation for *C. albicans* SC5314. Statistics were performed by Ordinary one-way Anova with Sidak’s multiple comparisons test. P values are shown as numeric values.
